# Intracellular *Salmonella* Paratyphi A is motile and differs in the expression of flagella-chemotaxis, SPI-1 and carbon utilization pathways in comparison to Intracellular *S*. Typhimurium

**DOI:** 10.1101/2021.08.18.456644

**Authors:** Helit Cohen, Claire Hoede, Felix Scharte, Charles Coluzzi, Emiliano Cohen, Inna Shomer, Ludovic Mallet, Remy Felix Serre, Thomas Schiex, Isabelle Virlogeux-Payant, Guntram Grassl, Michael Hensel, Hélène Chiapello, Ohad Gal-Mor

## Abstract

Although *Salmonella* Typhimurium (STM) and *Salmonella* Paratyphi A (SPA) belong to the same phylogenetic species, share large portion of their genome and express many common virulence factors, they differ vastly in their host specificity, the immune response they elicit, and the clinical manifestations they cause. In this work, we compared for the first time their intracellular trascriptomic architecture and cellular phenotypes during epithelial cell infection. While transcription induction of many metal transport systems, purines, biotin, PhoPQ and SPI-2 regulons was common in both intracellular SPA and STM, we identified 234 differentially expressed genes that showed distinct expression patterns in intracellular SPA vs. STM. Surprisingly, clear expression differences were found in SPI-1, motility and chemotaxis, and carbon (mainly citrate, galactonate and ethanolamine) utilization pathways, indicating that these pathways are regulated and possibly function differently, during their intracellular phase. Moreover, we show that the induction of flagella genes by intracellular SPA leads to cytosolic motility, a conserved trait specific to SPA. To the best of our knowledge, this is the first report of a flagellum-dependent intracellular motility of any *Salmonella* serovar in living host cells. Importantly, we demonstrate that the elevated expression of SPI-1 and motility genes by intracellular SPA results in increased invasiveness of SPA, following exit from host cells. We propose that such changes prime SPA towards new cycles of host cell infection and contribute to the ability of SPA to disseminate beyond the intestinal lamina propria of the human host, during enteric fever.

**IMPORTANCE:** *Salmonella enterica* is a ubiquitous, facultative intracellular animal and human pathogen. Although non-typhoidal *Salmonella* (NTS) and typhoidal *Salmonella* serovars belong to the same phylogenetic species and share many virulence factors, the disease they cause in humans is very different. While the underlying mechanisms for these differences are not fully understood, one possible reason expected to contribute to their different pathogenicity is a distinct expression pattern of genes involved in host-pathogen interactions. Here, we compared the global gene expression and the intracellular behavior, during epithelial cell infection of *S*. Paratyphi A (SPA) and *S*. Typhimurium (STM), as prototypical serovars of typhoidal and NTS, respectively. Interestingly, we identified different expression patterns in key virulence and metabolic pathways, together with intracellular motility and increased invasiveness of SPA, following exit from infected cells. We hypothesize that these differences are pivotal to the invasive and systemic disease developed following SPA infection in humans.

## INTRODUCTION

*Salmonella enterica* (*S. enterica*) is an abundant Gram-negative, facultative intracellular animal and human pathogen. This highly diverse bacterial species contains more than 2600 antigenically distinct serovars (biotypes) that are different in their host specificity and the disease they cause. The developed clinical manifestation is the result of multiple factors, but largely depends on the characteristics of the infecting serovar and the immunological status of the host (1). Many of the *S. enterica* serovars, including the ubiquitous *S. enterica* serovar Typhimurium (*S.* Typhimurium) are generalist pathogens and are capable of infecting a broad range of host species. Infection of immunocompetent humans by non-typhoidal *Salmonellae* (NTS) typically leads to a self-limiting, acute inflammatory gastroenteritis, confined to the terminal ileum and colon. In contrast, a few serovars including Typhi, Paratyphi A and Sendai, collectively referred to as ‘typhoidal salmonellae’ are restricted to the human host and known as the causative agents of enteric (typhoid) fever. In most cases, enteric fever is a non-inflammatory, systemic life-threatening disease, presented as bacteremia and dissemination of the pathogen to systemic sites such as the spleen, liver and lymph nodes (2–4).

*S. enterica* infections are still considered a significant cause of mortality and morbidity with an annual incidence of over 27 million cases of enteric fever (5), and 78.7 million cases of gastroenteritis (6) worldwide. In recent years, the global prevalence of *S.* Paratyphi A (SPA) is increasing and in some countries (especially in eastern and southern Asia), SPA infections are accountable for up to 50% of all enteric fever cases (7, 8). The lack of a commercial SPA vaccine and the increased occurrence of antibiotic resistant strains illuminate SPA as a significant public health concern that is still an understudied pathogen (9).

Active invasion into eukaryotic non-phagocytic cells is one of the key virulence-associated phenotypes of all *S. enterica* serovars. This unique capability facilitates *Salmonella* intestinal epithelium crossing of the small intestine (10) and is mediated by a designated type three secretion system (T3SS) encoded in the *Salmonella* Pathogenicity Island (SPI)-1. This sophisticated syringe-like nanosystem is evolutionary related to the flagellar apparatus (11) and used to translocate an array of effector proteins directly into the host cell cytoplasm. Translocated effectors by T3SS-1 trigger cytoskeletal rearrangements and *Salmonella* penetration of intestinal barriers (12). In addition, T3SS-1 and its associated effectors significantly contribute to intestinal inflammation (13) that helps *Salmonella* to compete with the gut microbiota (14).

After penetrating the lamina propria into the submucosa and mesenteric lymph nodes, *Salmonella* are taken up by phagocytic cells such as macrophages and dendritic cells. Within host cells, *Salmonella* are compartmentalized into a modified intracellular phagosome, known as the *Salmonella* containing vacuole (SCV). Inside the SCV *Salmonella* manipulates the phagosome-lysosome membrane fusion and other cellular pathways, and starts to replicate intracellularly (13). These activities require a second T3SS encoded by genes on SPI-2 and the translocation of a distinct set of effectors proteins (15).

Typhoid fever disease and host-response to *S*. Typhi infection is largely attributed to the function of the Vi polysaccharide capsule and its associated regulator, TviA encoded on SPI-7 (16–19). Nonetheless, since SPA does not harbor the SPI-7, nor expresses the Vi capsule, different mechanisms are expected to contribute to the clinically indistinguishable enteric fever disease caused by SPA and *S*. Typhi. Previously, we showed that in *Salmonella* culture grown in LB to the late logarithmic phase, SPI-1 genes and T3SS-1 effectors are expressed and secreted at significantly lower levels by SPA compared to *S*. Typhimurium (STM) (20), and demonstrated differences in the regulatory setup of the flagella-chemotaxis pathway between these serovars (21, 22).

Here, to further understand the different pathogenicity of SPA vs. STM as prototypic NTS, we set out to compare their transcriptional architecture, during non-phagocytic host cell infection. Surprisingly, in sharp contrast to the expression pattern in LB grown culture, we show that intracellular SPA is motile and induces the expression of SPI-1 and flagellar genes, compared to intracellular STM. In addition, we demonstrate that SPA and STM diverge in the intracellular expression of genes for citrate and ethanolamine metabolism and in their ability to utilize these carbon sources in vitro. We hypothesize that these differences are pivotal for the distinct diseases resulting from SPA vs. STM infection, and their different interactions with the human host.

## RESULTS

### The intracellular transcriptomic landscape of *S*. Typhimurium

Although STM and SPA belong to the same species, share about 89 % of their genes and express many common virulence factors (23), the disease they cause in immunocompetent humans is very different. One possible reason, expected to contribute to their different pathogenicity is a distinct expression of genes involved in host-pathogen interactions. To test this hypothesis, we compared the transcriptomic architecture and gene activity of SPA or STM in a disease-relevant context of epithelial cell infection. We chose to sample *Salmonella* infection at 8 h p.i. when the pathogen is considered to be well engaged in replication and optimally adapted to the intracellular environment (24). Thus, HeLa cells were infected with fluorescent STM and SPA for 8 h and sorted by flow cytometry to isolate *Salmonella*-infected cells. Isolated RNA was deep-sequenced (RNA-Seq), aligned to the corresponding *Salmonella* genome and compared to the transcriptome of *Salmonella* cultures growing extracellularly in rich LB medium.

RNA extracted from three independent infection experiments of HeLa cells infected with STM have generated 496-575 million RNA-Seq reads (human and bacterial reads) per experiment. Of which, 4.9-6.6 million sequence reads, assigned to non-rRNA bacterial genes were used for intracellular bacterial transcriptome analysis. Three independent extracellular STM cultures grown in LB, were used as a reference control, from which we obtained 4.3-13.8 million RNA-seq informative reads that were assigned to non-rRNA bacterial genes. To analyze the STM gene expression, we calculated for each gene the number of transcripts per kilobase million (TPM) from its feature read count counts. The expression level threshold was set to a TPM value of 10 or above (25). Altogether, we identified 1,018 distinct genes that were exclusively expressed by STM during intracellular infection of HeLa cells and 191 distinct genes that were expressed only when STM was grown in LB. 3,127 transcripts were commonly expressed at both conditions (Fig. 1A and Table S1).

**Figure 1.**
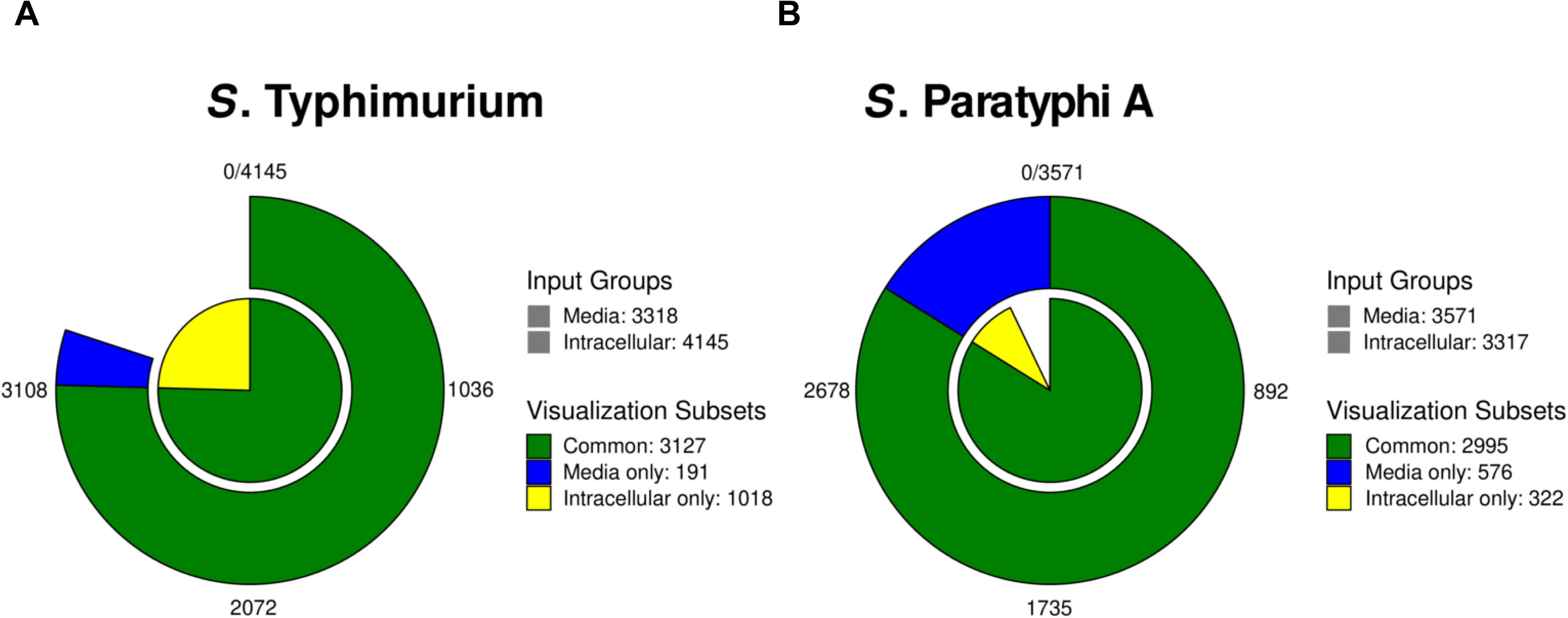
The number of STM and SPA genes expressed in LB medium and during intracellular growth. *Salmonella* RNA-seq reads were aligned to *S*. Typhimurium 14028S and *S*. Paratyphi A 45157 genomes and normalized by transcripts per million (TPM) transformation. Genes with TPM ≥ 10 were considered as expressed. The number of expressed genes in STM (**A**) and SPA (**B**) growing in LB medium to the late logarithmic phase and within HeLa cells at 8 h p.i. is shown.

RNA-seq successfully identified 365 upregulated and 465 downregulated genes that were changed by at least twofold inside epithelial cells at 8 h p.i. (adjusted p-value ≤ 0.05) by STM, relative to their expression in LB (Fig. 2A and Table S2). In agreement with previous studies (14, 26, 27), during STM infection, at least 27 genes belonging to the flagella-motility regulon and several genes (*fimAWYZ*) from the type I fimbriae were significantly repressed. Interestingly, the expression of genes encoding the Saf fimbriae (*safABCD*) were upregulated during HeLa cell infections by STM.

**Figure 2.**
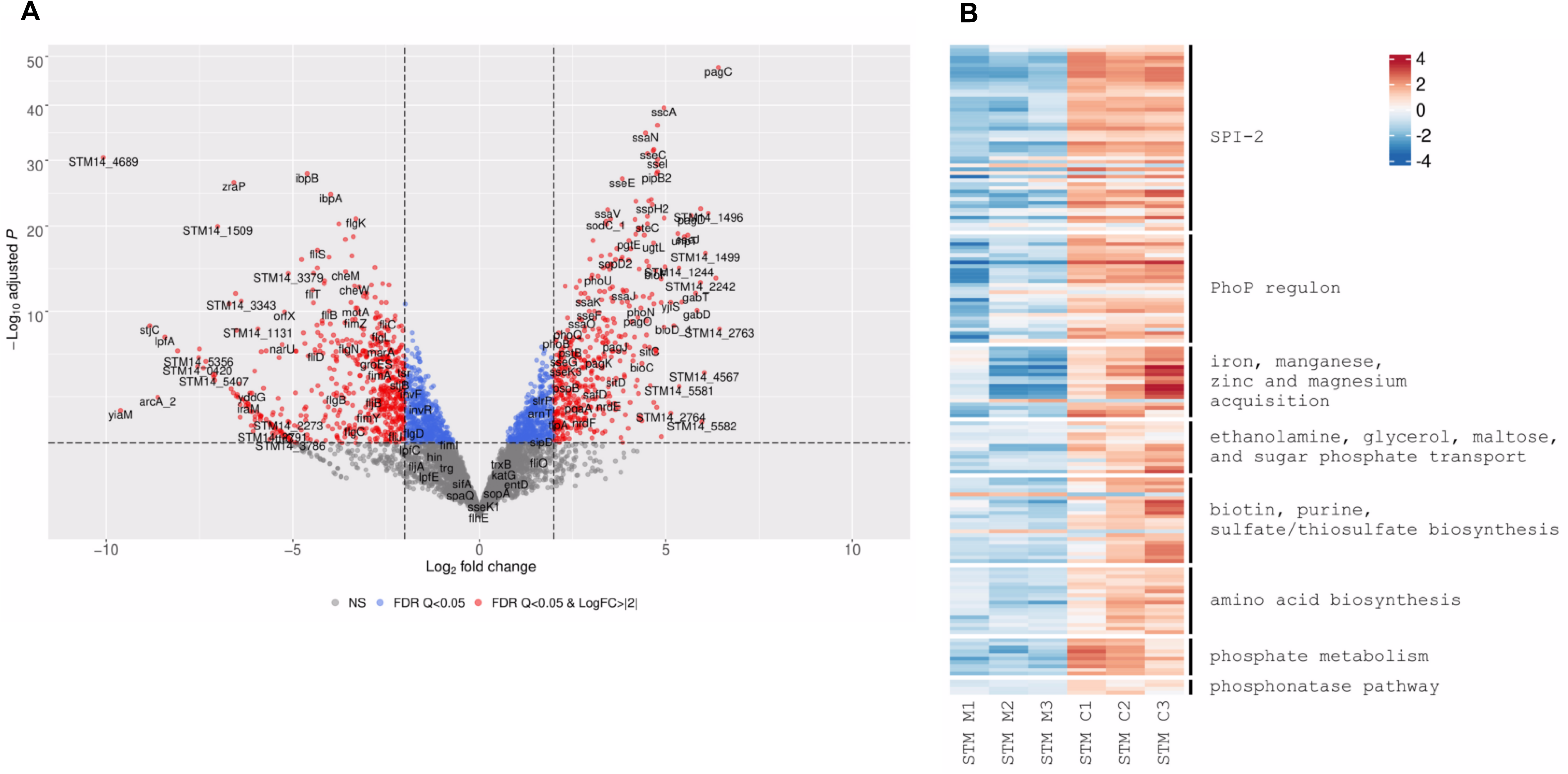
DEGs of intracellular *S*. Typhimurium compared to growth in LB medium. **(A)** Volcano plots showing the fold change (log2 ratio) in the expression of STM genes grown intracellularly in HeLa cells (8 h p.i.) vs. the growth in LB medium (X-axis) plotted against the -log_10_ adjusted p-value (Y-axis). Each dot on the plot represents the mean value (from three independent cultures) of one gene. Genes that were changed by more than twofold are colored in red. (**B**) Heatmap based on RNA-Seq results showing the relative transcription of STM genes of interest during intracellular infection (C1 to C3) vs. their extracellular expression in LB medium (M1 to M3).

In contrast to the flagella and the *fim* genes, at least 31 genes from the PhoPQ regulon were highly induced intracellularly (Fig. 2B). The two-component system PhoPQ, orchestrates *Salmonella* adaptation to the intracellular milieu and regulates a wide array of genes central for *Salmonella* virulence including those encoded on SPI-2 (28, 29). Correspondingly, we identified upregulation of the SPI-2 regulon including *ssaBCDEGHIJKLMVNOPRSTU*, *sseABCDEFGIJL*, *sseK3*, *sspH2*, *pipB, pipB2*, and *steC* genes (Fig. 2B). These results are consistent with the known notion that SPI-2 expression is induced during intracellular replication of *Salmonella* (15, 30) and with previous reports that studied STM expression during macrophages infection (31, 32). Other PhoP-regulated genes including *pgtE* (involved in bacterial resistance to antimicrobial peptides), *pagN* (encoding an adhesion/invasion protein), *phoN* (acid phosphatase) and *pagCDJKO* were also induced (Fig. 2B).

Iron is an essential micronutrient required by nearly all bacterial species, including *Salmonella.* Limiting the availability of the nutrient metals to intracellular pathogens is one of the main mechanisms of nutritional immunity. In return, *Salmonella* uses a variety of high-affinity iron uptake systems to compete with the host for essential transition metals required for its intracellular growth (33). Indeed, we identified elevated expression of multiple iron (*entABEFH* and *iroNBCDE*), iron/ manganese (*sitABCD*), zinc (*zntR*, *zitT* and *znuA*), and magnesium (*mgtAB*) acquisition system genes, highlighting the essentiality of these metal ions for STM intracellular replication (Fig. 2B).

More than 100 suitable carbon substrates as well as various nitrogen, phosphorus and sulfur sources are available to invading pathogens, at different niches of the vertebrate hosts (34, 35). Interestingly, multiple genes belonging to the ethanolamine operon (*eutQTMGAC*) and genes involved in glycerol (*glpABDK*), and maltose (*malEKM*) metabolism were particularly induced during STM infection (Fig. 2B). Nevertheless, the most highly induced metabolic gene was *uhpT,* encoding a hexose phosphate transporter that was upregulated by more than 40-fold at 8 h p.i. These results suggest that sugar phosphates are major substrates required for cytosolic intracellular proliferation of STM within HeLa cells, but also that ethanolamine, glycerol and maltose could be utilized as alternative carbon sources in epithelial cells.

Similarly, biotin (*bioABCDF*), purine (*nrdEFHI* and *purDEHKLT*), and sulfate/thiosulfate (*cysACHITW* and *sufABCDES*) biosynthetic metabolic pathways were highly transcribed intracellularly by STM. Moreover, genes involved in biosynthesis of the amino acids histidine (*hisACDFH* and *hutG*), tryptophan (*trpABDE*), and glutamine (*glnAGKL*), and genes that catalyze the cleavage of N-acetylneuraminic acid (sialic acid; Neu5Ac) to form pyruvate and N-acetylmannosamine (ManNAc) (*nanAEKT* and *nagB*) were also upregulated by STM during HeLa cell infections (Fig. 2B).

A large number of genes involved in phosphate metabolism were identified to be induced within HeLa cells. For example, the transcriptional levels of *pstABCS* (encoding an ATP-dependent phosphate uptake system), which is responsible for inorganic phosphate uptake during phosphate starvation, *ugpABCEQ, phoR* and *apeE* were markedly increased. The *ugp* operon is involved in hydrolysis of diesters during their transport at the cytoplasmic side of the inner membrane (36). Thereby, sn-glycerol-3-phosphate (G3P) is generated and used as a phosphate source. The *upg* operon is regulated by a two-component system, PhoBR, which also regulates the outer membrane esterase, ApeE (37). Additional genes belonging to the PhoB regulon, such as *phnSVWU* encoding the phosphonatase pathway, which is responsible for breaking down phosphonate and yielding cellular phosphate (38) were also significantly induced.

Collectively, these results suggest that STM likely experiences iron, biotin, purine and phosphate deprivation in host epithelial cells and induces the expression of genes involved in their transport and utilization. In addition, these results imply that ethanolamine, glycerol and maltose are available sources of carbon for STM in epithelial cells.

### The intracellular transcriptomic landscape of *S*. Paratyphi A

RNA-Seq analysis was similarly applied for intracellular SPA. 313.8 to 805.9 million dual-RNASeq reads were obtained from three independent HeLa cell cultures infected with SPA. Out of these, 14.6 to 36.6 million sequence reads were assigned to non-rRNA SPA genes and used to determine the intracellular SPA transcriptome (Table S3). As a reference control, RNA extracted from SPA grown in LB from three independent cultures generated 7.3 to 23 million informative reads (assigned to non rRNA SPA genes). Overall, we identified 322 genes that were exclusively expressed (TPM ≥10) by SPA during infection of HeLa cells compared to 576 genes that were expressed only by extracellular SPA in LB culture. 2,995 SPA genes were expressed at both conditions (Fig. 1B and Table S3).

At 8 h p.i. of HeLa cells, 322 genes were upregulated and 579 genes were down-regulated (Fig. 3A and Table S4) by twofold or more (adjusted p-value ≤ 0.05). Like in intracellular STM, the genes for type I (Fim) fimbriae (*fimCIWYZ*) and additional fimbria clusters including Bcf (*bcfDE*), the Curli (*csgBCDE*), Stb (*stbBE*), Stc (*stcABC*), Stf (*stfAD*), Sth (*sthABCD*), and Stk (*stkABC*) were all strongly downregulated intracellularly (Fig. 3B). Noteworthy, the downregulation of type I (*fimACIZ*), Curli (*csgBDE*) and Bcf (STM14_0037 and *bcfB*) fimbriae was more pronounced in SPA than in STM (Fig. S1), indicating that the SPA fimbriome is robustly repressed, during intracellular replication, while some basal level of expression of these fimbrial genes is still maintained during STM infection.

**Figure 3.**
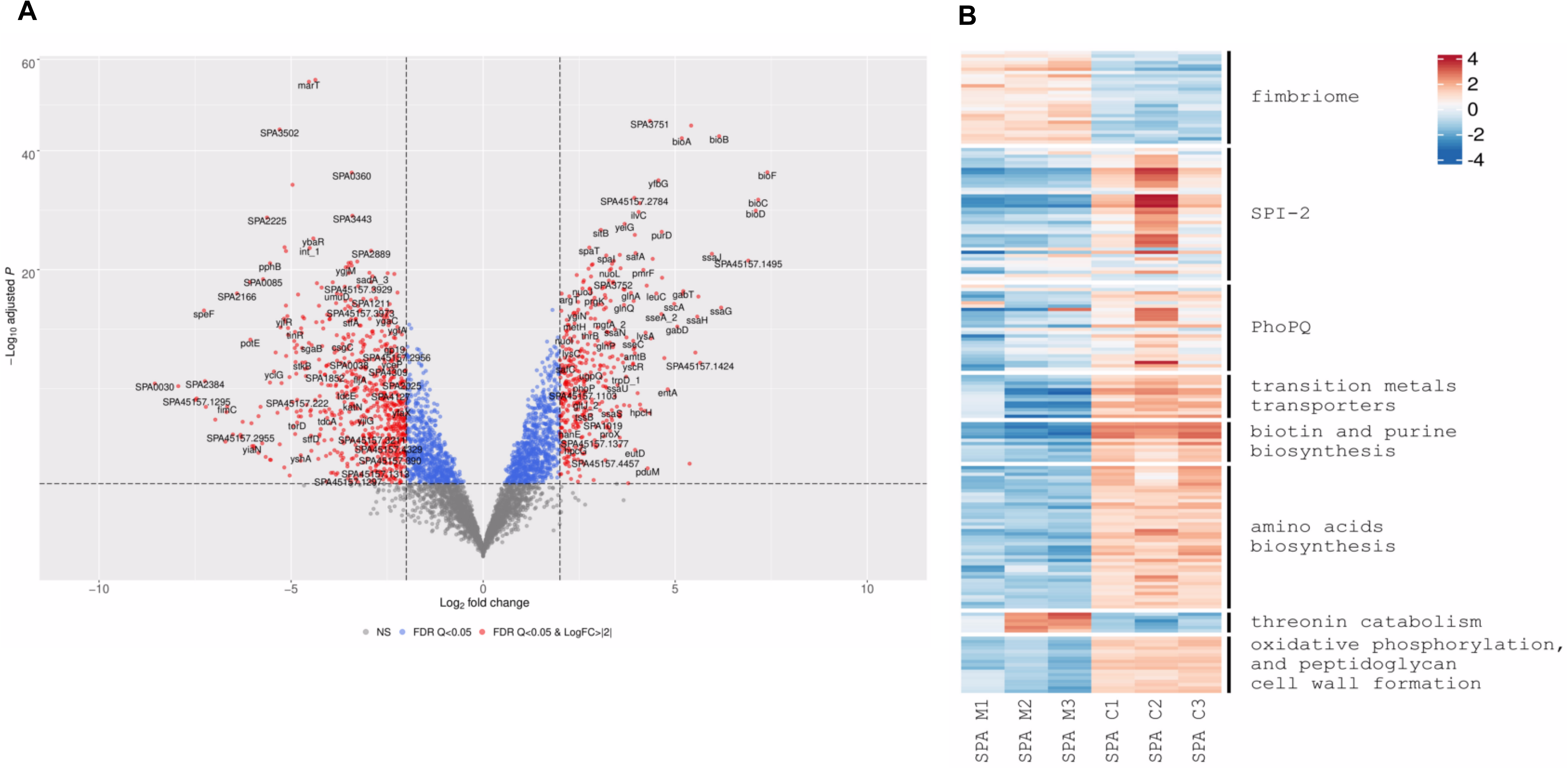
DEGs of intracellular *S*. Paratyphi A compared to growth in LB medium. **(A)** Volcano plots showing the fold change (log2 ratio) in the expression of SPA genes grown intracellularly in HeLa cells (8 h p.i.) vs. the growth in LB medium (X-axis) plotted against the -log_10_ adjusted p-value (Y-axis). Each dot on the plot represents the mean value (from three independent cultures) of one gene. Genes that were changed by more than twofold are colored in red. (**B**) Heatmap based on RNA-Seq results showing the relative transcription of SPA genes of interest during intracellular infection (C1 to C3) vs. their extracellular expression in LB medium (M1 to M3).

In contrast, but consistent with our findings for intracellular STM, genes belonging to the PhoPQ (including polymyxins resistance) and SPI-2 regulons were significantly induced by intracellular SPA. Moreover, various transition metals import systems including manganese and iron transporter (*sitABCD*), enterobactin biosynthesis system (*entABCDEF* and *fepAG*) and the iron ABC transporter IroC were considerably upregulated during SPA HeLa cell infection (Fig. 3B).

In addition, the expression of several metabolic pathways that were significantly elevated during STM infection were induced by intracellular SPA as well, including biotin (*bioABCDF*) and purine (*purCDEHKLT*) biosynthesis genes. Further overlap in the intracellular transcriptome of these pathogens includes several amino acid biosynthesis pathways such as glutamine (*glnAGKLPQ*), tryptophan (*trpABCDE*) and histidine (*hisAFGHI*) that were upregulated in both intracellular STM and SPA.

Nevertheless, comparison of the intracellular transcriptome between STM and SPA, during HeLa cells infection, identified 234 differentially expressed genes (DEGs; Table S5), indicating that some pathways may function differently during the intracellular phase of STM vs. SPA. These differences included the induction of several amino acids biosynthesis genes which expression was more pronounced in intracellular SPA than in STM. These include biosynthesis genes of glycine (*glyASU*), isoleucine (*ilvCDEG*), leucine (*leuABCD*), lysine (*lysAC* and *sucBD*), methionine (*metCEFHKLR*), serine (*serAC*), and threonine (*thrBC*). In contrast, the threonine degradation pathway (*tdcABCDEG*) was found to be repressed in intracellular SPA. These results may suggest that SPA relies more than STM on amino acids as an available carbon and energy source during epithelial cell infection. Additional pathways that were found upregulated in SPA, but not in STM are the oxidative phosphorylation (*nuoEFGHIJKLMN* and *atpCD*), and peptidoglycan cell wall formation (*murACDEG*) pathway genes (Fig. 3B). Remarkably, further differences between the intracellular transcriptome of SPA vs. STM included different expression profiles of the SPI-1, motility and chemotaxis, and carbon utilization regulons as detailed in the following sections.

### Differences in carbon catabolism during intracellular infection of *S*. Paratyphi A and *S*. Typhimurium

In order to survive, persist and replicate in host cells, intracellular pathogens must adapt their metabolic pathways to the specific physical conditions and available nutrients found in the intracellular environment. Carbon catabolism provides bacteria with energy by means of reducing equivalents, ATP and essential biosynthetic precursors. Different studies have suggested that hexose monosaccharides such as glucose and glucose-6P are the main source of carbon during *Salmonella* infection (39–43). Comparison of the metabolic gene expression between intracellular STM and SPA revealed significant differences in the expression profile of several carbon catabolic pathways. For example, the gene *uhpT* encoding an antiporter for external hexose 6-phosphate and internal inorganic phosphate (44) presented about 90-fold higher expression by intracellular STM than SPA (Fig. 4A). This different expression might be because the sensor-regulator system encoded by *uhpABC* is defective by pseudogene formation in SPA (45). Similarly, the gene coding for galactonate permeases (*dgoT*) was upregulated in intracellular STM, but not in SPA, suggesting that STM utilizes more hexose 6-phosphate and galactonate than SPA as a carbon source during intracellular growth.

**Figure 4.**
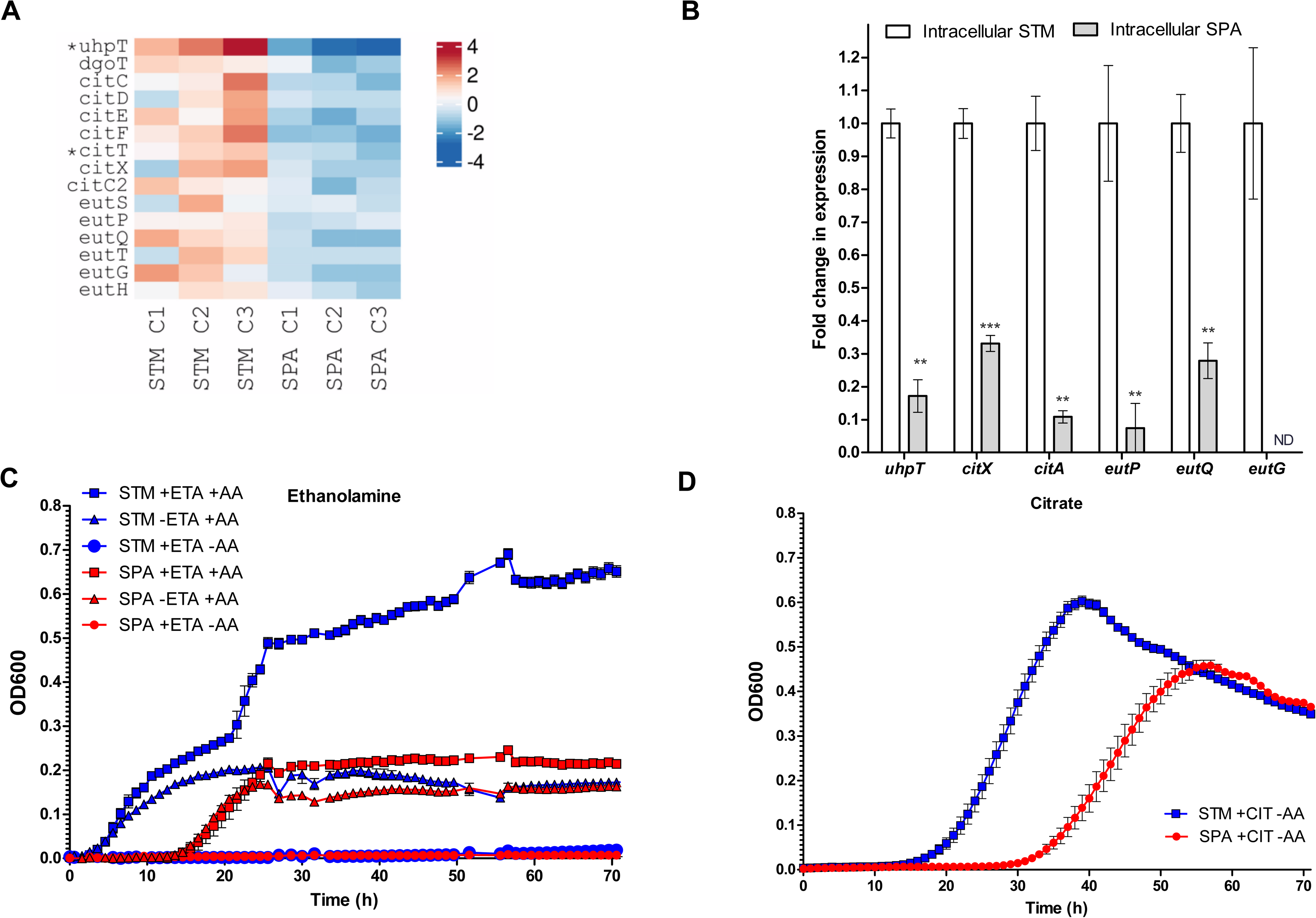
Differences in carbon utilization gene expression between STM and SPA. (**A**) Heatmap of carbon catabolism gene expression by intracellular STM vs. intracellular SPA. Pseudogenes in the SPA genome are marked with an asterisk. (**B**) The fold change in the transcription of the glucose-6P (*uhpT*), citrate (*citA* and *citX*), and ethanolamine (*eutP, eutQ* and *eutG*) utilization pathways in intracellular STM relative to their expression in intracellular SPA was determined using qRT-PCR. RNA was extracted from FACS-sorted intracellular salmonellae 8 h p.i. The mean and the standard error of the mean of 3-8 independent qRT-PCR reactions are shown. Student t-test was used to determine statistical significance. **, p < 0.005; ***, p < 0.0001; ND, not detected. (**C**) Growth of STM and SPA in M9 in the presence (+) or absence (-) of casamino acids (AA) and ethanolamine (ETA), under anaerobic conditions. (**D**) Growth of STM and SPA in M9 supplemented with citrate as a sole carbon source.

Similarly, RNA-Seq identified that at least seven genes involved in citrate metabolism including *citCDEFTX* and *citC2* exhibited higher transcription levels in intracellular STM than SPA (Fig. 4A). Independent qRT-PCR further confirmed that the intracellular expression of *citA* and *citX* encoding citrate-proton symporter and apo-citrate lyase phosphoribosyl-dephospho-CoA transferase, respectively is lower by about 3- and 10-fold in SPA infected cells vs. STM (Fig. 4B).

STM can utilize ethanolamine as a sole source of carbon, nitrogen, and energy in a cobalamin (vitamin B12)-dependent manner. Ethanolamine may be important to STM growth in the host, since it is derived from the membrane phospholipid phosphatidylethanolamine that is particularly prevalent in the gastrointestinal tract. This process involves breakdown of the precursor molecule phosphatidylethanolamine by phosphodiesterases to glycerol and ethanolamine (46). In STM, ethanolamine catabolism involves 17 genes organized in the *eut* operon, controlled by the transcriptional activator EutR (47). At least six ethanolamine catabolism genes (*eutSPQTGH*) were expressed at higher levels by intracellular STM compared to SPA (Fig. 4A). These differences were independently verified by qRT-PCR that showed significantly reduced expression of *eutG, eutP* and *eutQ* genes in intracellular SPA vs. STM (Fig. 4B).

Collectively, these observations suggest that STM is able to better utilize intracellular citrate, ethanolamine and galactonate as intracellular carbon sources than SPA, and that SPA may use an alternative carbon source such as amino acids during intracellular growth.

To further test this hypothesis, we compared the growth of SPA and STM in a defined M9 medium supplemented with ethanolamine or citrate as a carbon source. Growth under anaerobic conditions in M9 medium, supplemented with ethanolamine, without amino acids, did not allow bacterial growth of neither STM nor SPA. Adding casamino acids, without additional carbon source, resulted in a minimal and comparable growth of both STM and SPA, likely due to the utilization of amino acids as a restricted carbon source. Nevertheless, adding ethanolamine as a *bona fide* carbon source to the medium resulted with increased and rapid growth of STM, but with a minor improvement of SPA growth (relative to its growth on amino acids only; Fig. 4C), indicating the ability of STM, but not SPA to efficiently ferment ethanolamine. Replacing the carbon source with citrate under aerobic growth conditions allowed delayed, but substantial growth of STM, in comparison to restricted and slower growth of SPA (Fig. 4D). These results are concurring with the expression data and indicate a better ability of STM than SPA to utilize both ethanolamine and citrate as a sole carbon source. These differences may also contribute to a higher intracellular replication pace of STM than SPA found in HeLa cells (Fig. S2).

### Intracellular *S*. Paratyphi A expresses SPI-1 genes at higher levels than intracellular *S*. Typhimurium

Previously we showed that under extracellular aerobic growth conditions in LB, SPI-1 gene expression, as well as secretion of SPI-1-T3SS effector proteins, occur at significantly lower levels in SPA compared to STM (20). Here, in sharp contrast, we found that intracellular SPA transcribed significantly higher levels of SPI-1 and T3SS-1-effector genes than intracellular STM (Fig. 5A). This was clearly evident (log 2 change ≥ 2 and p-value ≤ 0.05) for at least 14 SPI-1 genes including the SPI-1 regulators *hilA* and *hilD*, the effector genes *sopBDF*, the T3SS-1 structural genes *invCHIJ* and *spaOPQ*, *sigE* encoding a chaperon (a.k.a. *pipC*) and *iagB* (Table S5).

**Figure 5.**
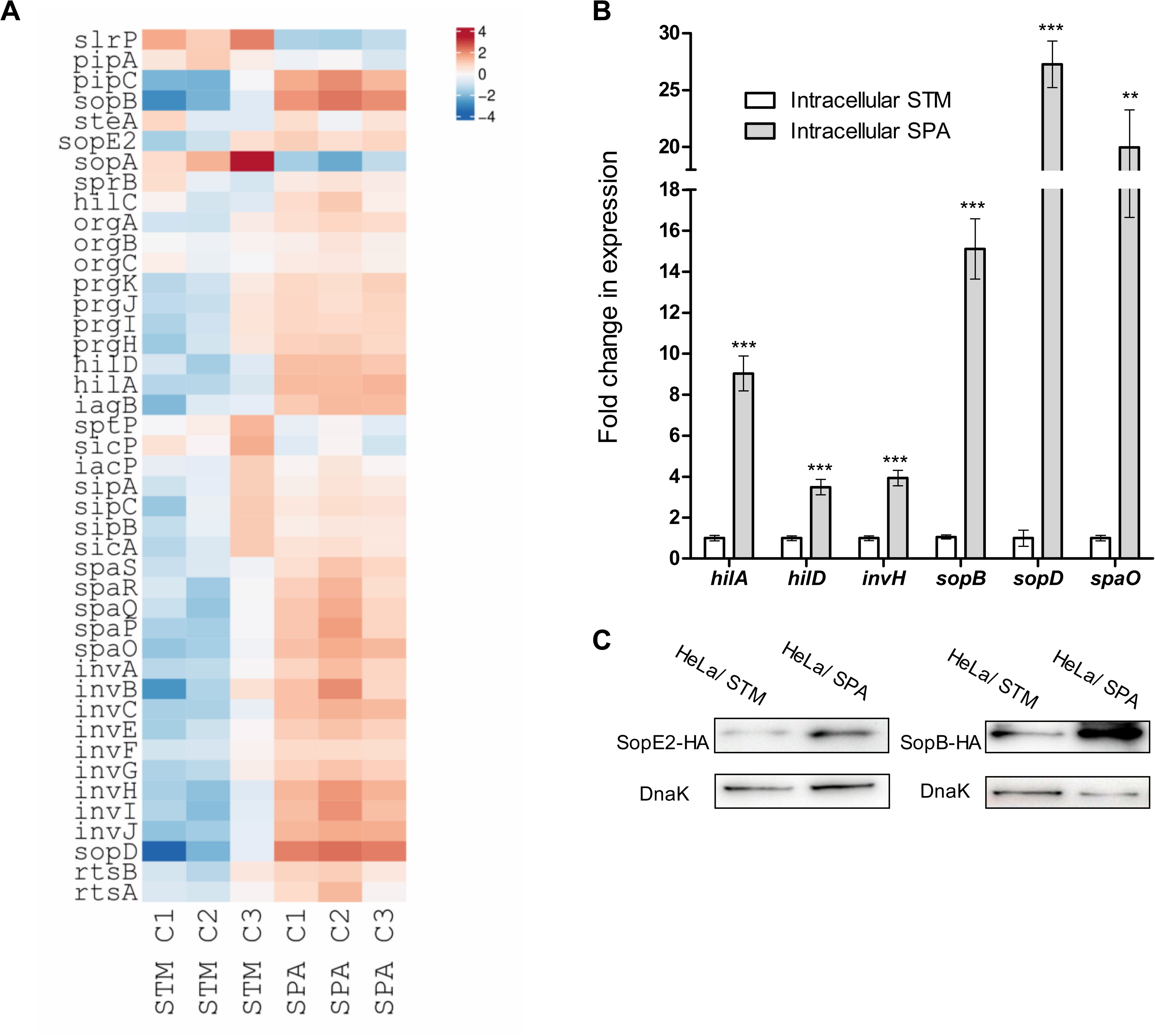
Intracellular SPA expresses higher levels of SPI-1 genes than intracellular STM. (**A**) Heat map of RNA-Seq results showing the relative transcription of SPI-1 genes in three independent HeLa cell infections with STM and SPA. (**B**) The fold change in the transcription of six SPI-1 genes (*hilA, hilD, invH, sopB, sopD* and *spaO*) in intracellular SPA relative to their expression in intracellular STM was determined by qRT-PCR. The results show the mean of 3-6 qRT-PCR reactions and the error bars indicate the SEM. (**C**) SPA and STM strains expressing GFP and a 2HA-tagged version of the SPI-1 effectors SopE2 and SopB were used to infect HeLa cells. At 8 h p.i., cells were fixed and FACS-sorted for infected (GFP-positive cells). Infected HeLa cells (1×10^6^ for each sample) were collected and separated on 12% SDS-PAGE. Western blotting using anti-hemagglutinin (HA) tag antibodies was used to detect the intracellular expression of SopE2 and SopB. Anti-DnaK antibodies were used as a loading control.

Independent qRT-PCR was used to verify these observations and demonstrated that intracellular SPA transcribed *hilA*, *hilD*, *invH*, *sopB*, *sopD* and *spaO* at 4- to 27-fold higher levels than STM (Fig. 5B). Moreover, Western blotting against a 2HA-tagged version of SopB and SopE2 further indicated elevated translation of these two SPI-1 effectors in intracellular SPA vs. STM (Fig. 5C). Collectively these results indicate that SPA expresses SPI-1 genes at higher levels during infection of non-phagocytic cells, in comparison to intracellular STM.

### Intracellular *S*. Paratyphi A is motile and expresses flagella-chemotaxis genes at higher levels than *S*. Typhimurium

Functional flagella are necessary for *Salmonella* motility and invasion of host cells (21, 48, 49). In *S. enterica,* more than 50 genes are involved in the assembly and motion of flagella, organized in one regulon that composes three ordered regulatory phases corresponding to classes I (early), II (middle), and III (late) (50).

Previous studies have shown that the genes coding the flagellar machinery, as well as many genes involved in chemotaxis, are strongly repressed during STM intracellular lifestyle (27, 31, 32). Here, RNA-Seq analysis has demonstrated that at least 38 out of 50 chemotaxis and flagella genes, belonging to class II and III of the flagella regulon are expressed at significantly higher levels in intracellular SPA compared to STM (Fig. 6A and Table S5). These include the flagellin gene *fliC* that was expressed at 11-fold higher levels by SPA vs. STM, and many other flagellar genes including *flgC* that demonstrated 64-fold difference in its expression between intracellular SPA vs. STM. In addition to the flagellar genes, SPA residing in the host cell presented much higher expression levels of chemotaxis genes (e.g. *cheB, cheR, cheM, cheW, cheZ*, and *cheY*) than intracellular STM.

**Figure 6.**
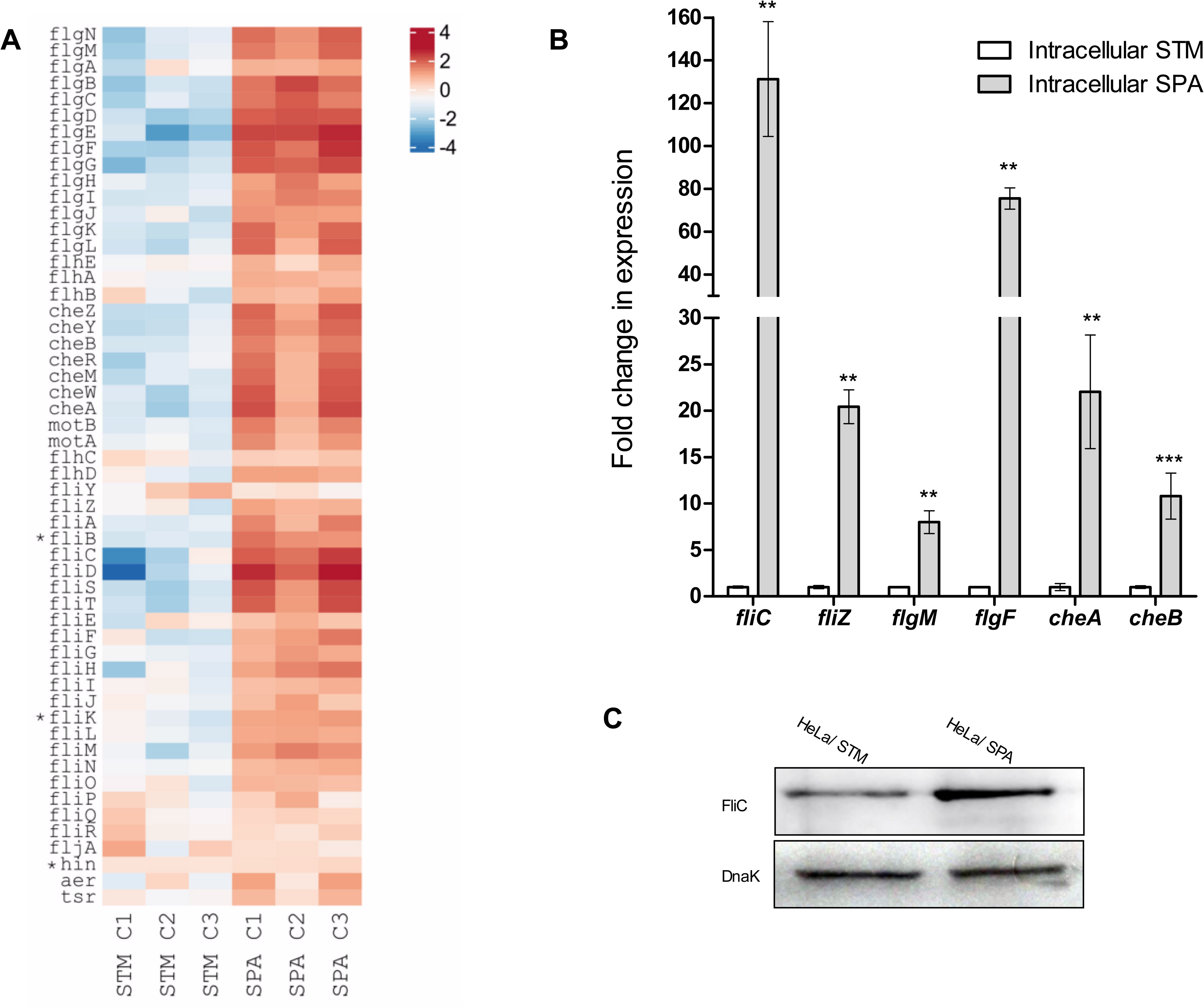
Intracellular SPA expresses higher levels of motility-chemotaxis genes than intracellular STM. (**A**) Heat map of RNA-Seq results showing the relative transcription of the motility-chemotaxis regulon in three independent HeLa cells infections with STM and SPA. (**B**) The fold change in the transcription of six flagellum genes (*fliC, fliZ, glgM, flgF, cheA* and *cheB*) in intracellular SPA relative to their expression in intracellular STM was analyzed by qRT-PCR. The results show the mean of 3-5 independent reactions and the error bars indicate the SEM. (**C**) SPA and STM strains expressing GFP were used to infect HeLa cells. At 8 h p.i., cells were fixed and FACS-sorted. Equal numbers of 1×10^6^ infected HeLa cells were collected, lysed, and proteins were separated on 12% SDS-PAGE. Western blotting using anti-FliC antibodies were used to detect synthesis of FliC by intracellular *Salmonella*. Anti-DnaK antibodies were used as a loading control.

Independent qRT-PCR analysis confirmed these results and showed that in sorted infected HeLa cells, the expression of *fliC, fliZ, flgM, flgF, cheA* or *cheB* was 8- to 130-fold higher in intracellular SPA than in intracellular STM (Fig. 6B). In agreement with the RNA-seq and qRT-PCR analyses, Western blotting using anti-FliC antibodies demonstrated significantly higher translation of FliC in intracellular SPA vs. STM (Fig. 6C). Taken together, these findings uncover significantly higher expression levels of motility and chemotaxis genes by SPA vs. STM within infected HeLa cells.

To further analyze if the increased expression of motility genes by intracellular SPA has physiological consequences, we investigated the possible presence of flagella in intracellular SPA. Immunostaining of intracellular SPA at 8 h p.i. indeed indicated the presence of flagella filaments on intracellular SPA (Fig. S3). Flagella staining was absent for a Δ*fliC* strain of SPA, and restored in the plasmid-complemented SPA Δ*fliC*. No immunolabeling of flagella was observed for intracellular STM WT or STM Δ*fliC* Δ*fljB* (Fig. S3).

### Motile intracellular SPA are more inclined to replicate in the cytoplasm rather than the SCV

Analysis of host cells infected by STM or SPA WT or T3SS-2-deficient strains was performed. To distinguish *Salmonella* in the SCV from *Salmonella* in other compartments or in the cytosol, and to enable live-cell imaging of infected host cells, we used LAMP1-GFP expressing HeLa host cells (Fig. 7, Movies 1, 4). Lifeact-GFP expressing HeLa cells were used to investigate potential interactions with the host cell actin cytoskeleton (Movie 1-3).

**Figure 7.**
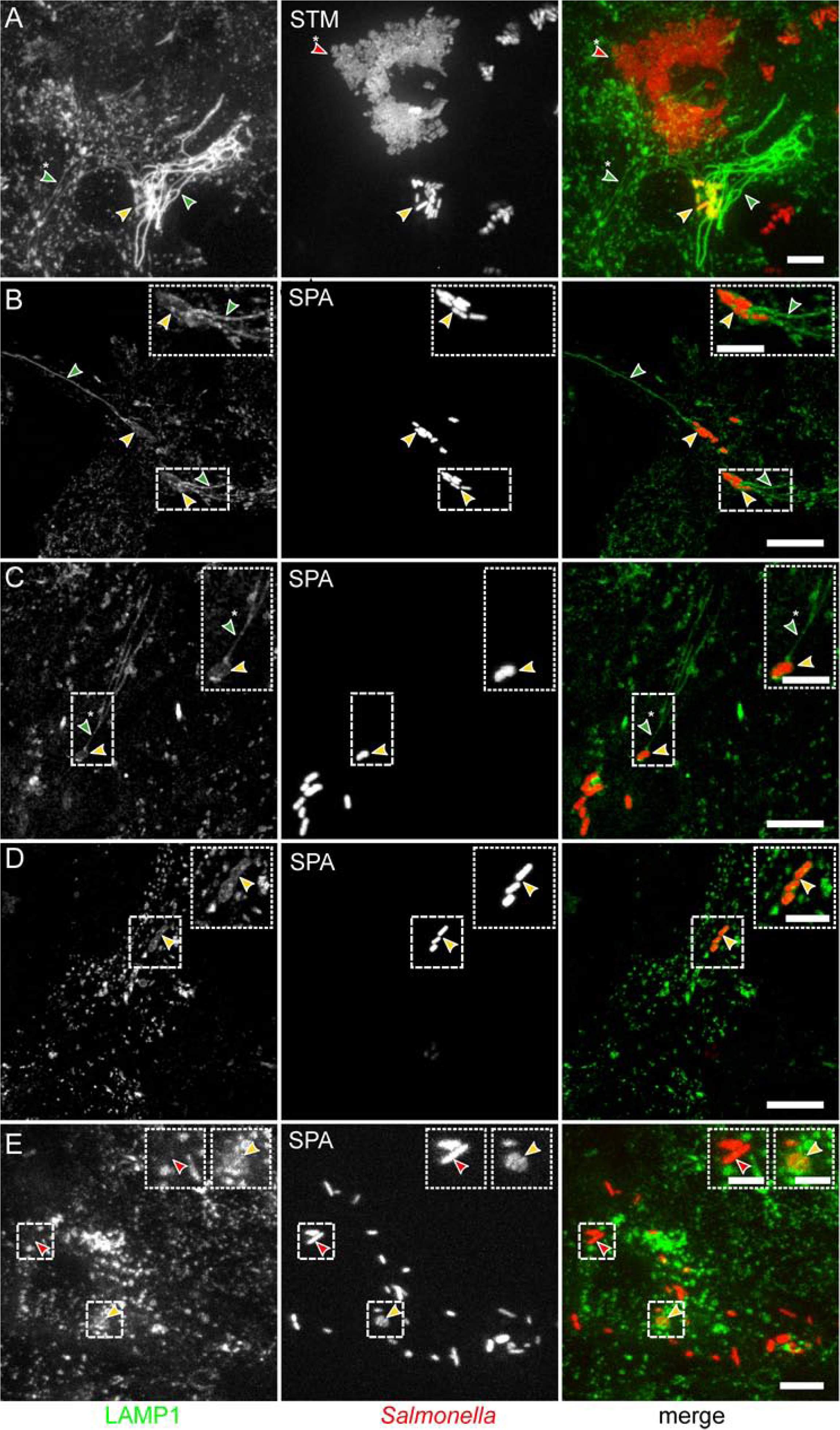
Distinct phenotypes of intracellular SPA. HeLa cells stably transfected for expression of LAMP1-GFP (green) were infected with STM WT (**A**, MOI 25) or SPA WT (**B-E**, MOI 75), each harboring pFPV-mCherry for constitutive expression of mCherry (red). Live-cell imaging was performed at 8 h p.i. Images are shown as maximum intensity projection. Arrowheads indicate subpopulations with distinct phenotypes in infected cells: SIF (green), thin SIF (green with asterisk), *Salmonella* residing in SCV (yellow), cytosolic *Salmonella* (red), hyper-replicating cytosolic *Salmonella* (red with asterisk). Scale bars: 10 µm (overview), 5 µm (inset).

More than 80% of STM WT were present in SCV with associated *Salmonella*-induced filaments (SIFs; Fig. 7A, Fig.8). STM WT was observed residing in SCV and the majority of infected host cells showed formation of extensive SIF networks. The STM *ssaV* strain was typically present in SCV, but tubular endosomal filaments were almost absent. A smaller proportion of host cells (ca. 10%) harbored cytosolic STM, and for these cells, massive cytosolic proliferation or hyper-replication was evident (Fig. 7A). For intracellular SPA WT, more heterogeneous phenotypes were observed (Fig. 7BCDE). SPA was located residing in the SCV, or present in the cytosol without association with LAMP1-positive membranes. Intracellular SPA also induced SIF in a T3SS-2-dependent manner, but the frequency of SIF-positive cells was much lower with ca. 9% for SPA WT compared to more than 80% for STM WT (Fig. 7BC, Fig. 8). SPA-infected host cells often contained thin SIF, that are likely composed of single-membrane tubules, in contrast to SIF with more intense LAMP1-GFP signal, indicating double-membrane tubules (Fig. 7BC). For SPA WT, SIF formation was significantly less frequent, yet dependent on T3SS-2. The frequency of SPA associated with SCV marker LAMP1 was highly reduced compared to STM. As intracellular SPA was observed to reside in tightly enclosing SCV (51), we considered flagella expression by vacuolar SPA less likely than by cytosolic bacteria. Because transcriptional analyses showed expression of flagellar genes by intracellular SPA, flagella filaments were detected on intracellular SPA, and a major proportion of infected HeLa cells (∼ 40%) harbored cytosolic SPA, we investigated the functional consequences of flagella expression by live-cell imaging of *Salmonella*-infected host cells (Movies 1-4). Indeed, we observed actively motile intracellular SPA in about 40% of the infection host cells.

**Figure 8.**
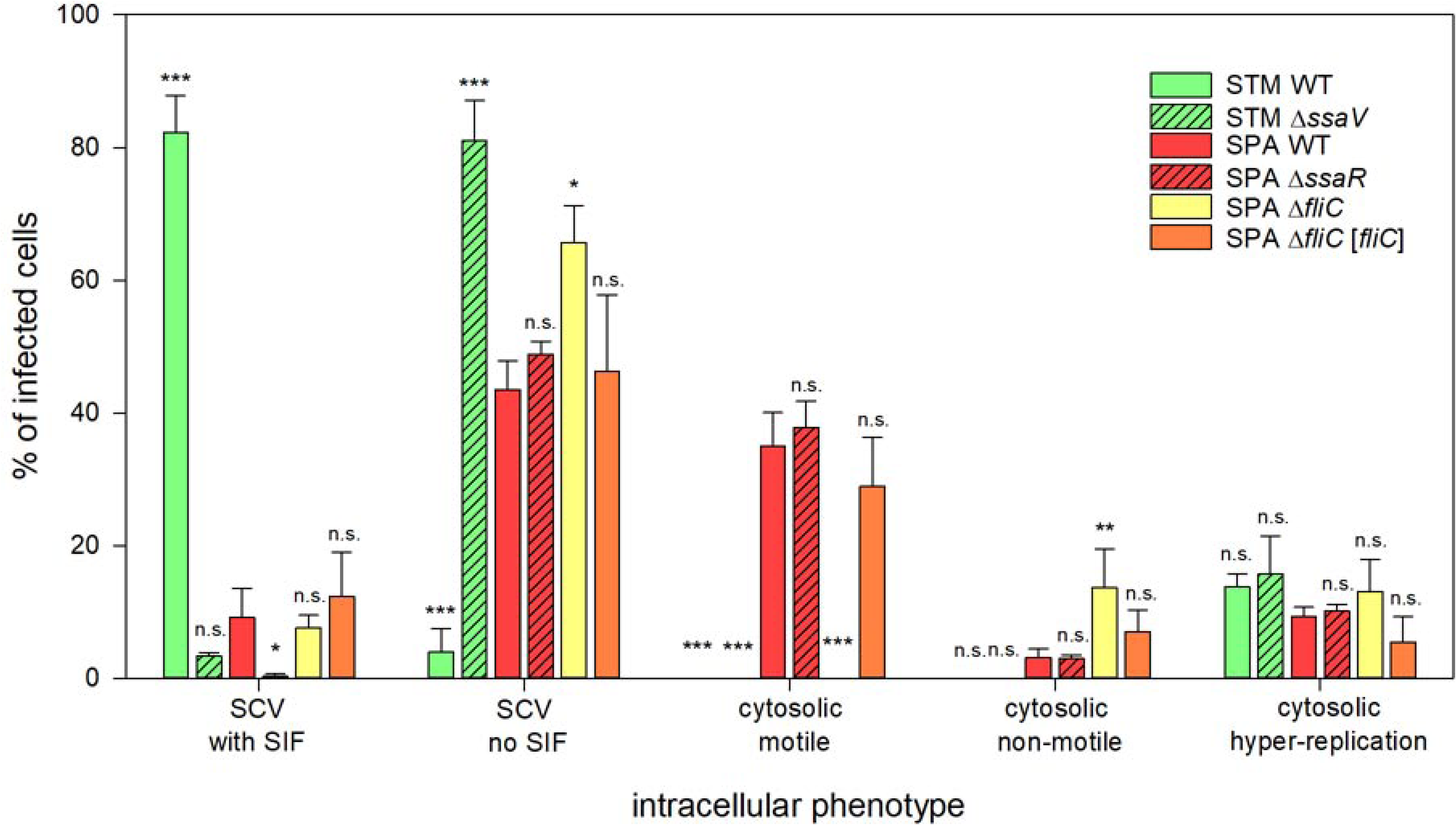
Quantification of intracellular phenotypes. HeLa cells stably expressing LAMP1-GFP (green) were infected with *Salmonella* strains expressing mCherry (red). Quantification of intracellular phenotypes was performed with a Zeiss Cell Observer with 63x objective. From 6-8 h p.i. at least 100 infected cells per strain (for SPA *fliC* at least 50) were examined for intracellular phenotype. Data represent the means with standard deviations for three biological replicates for each strain and condition. Statistical analysis was performed with One-way ANOVA and is indicated as n.s., not significant; *, p < 0.05; **, p < 0.01; ***, p < 0.001.

As motility in intracellular microbes typically is mediated by polymerization of host cell actin by microbial surface proteins (52), we investigated F-actin dynamics in SPA-infected HeLa Lifeact-GFP cells (Movie 1). No association of F-actin with static or motile intracellular SPA was evident, and F-actin trails as observed for intracellular motile *Listeria monocytogenes* (53) were absent in SPA WT infected cells. No cytosolic motility was observed for intracellular STM. To control if cytosolic motility is linked to the observed flagella expression, we used a SPA *fliC* mutant strain. This non-motile mutant completely lacked cytosolic motility, but showed increased presence in SCV (Fig. 8, Movie 1). Cytosolic motility of SPA was not affected in the T3SS-deficient Δ*ssaR* strain (Fig. 8, Movie 1). Live-cell imaging at high frame rates (Movies 2 and 3) demonstrated the dynamics of intracellular motility of SPA in real time, with characteristic alternations between straight swimming and stops with tumbling. Analyses of the vectors of individual SPA indicated collisions with the host cell plasma membrane, but there were no indications for formations of protrusions as observed for intracellular motile *L. monocytogenes* (53).

We next set out to analyze the broader relevance of intracellular motility of SPA and investigated a set of clinical SPA isolates (22), and SPA reference strains of SARB collection (54). We observed that all SPA strains tested showed intracellular motility in HeLa cells (Movie 4). Expression of flagella genes and presence of flagella filaments was previously demonstrated for STM in dead host cells extruded from polarized epithelial monolayers (55). As these experiments were performed with STM strain SL1344, we also analyzed this strain in our experimental setting. As for STM ATCC 14028, also STM SL1344 showed cytosolic hyper-replication in infected HeLa cells, but cytosolic motility was absent (Movie 4).

These data confirm that the expression of flagella by intracellular SPA leads to flagella-mediated cytosolic motility, and that this motility is a conserved trait specific to SPA.

### Intracellular SPA are primed for reinfection of host cells

We set out to investigate potential functional consequences of the increased expression of SPI-1 and motility genes by intracellular SPA. We hypothesized that increased expression of SPI-1, higher amounts of T3SS-1 effector proteins, and motility could affect the interaction of SPA released from infected cells with naïve host cells. As host cell decay and release of the intracellular *Salmonella* is asynchronous, and quantification of new infection events is difficult in gentamicin protection assays, we set up a re-invasion assay that provides experimental control. HeLa cells were infected by invasive *Salmonella*, at a defined time point of 8 h p.i., host cells were osmotically lysed, and released bacteria were used to reinfect new epithelial cells. The number of viable bacteria in inoculum, released from initially infected host cells, and the number of bacteria in the second round of invasion were quantified (Fig. 9A).

**Figure 9.**
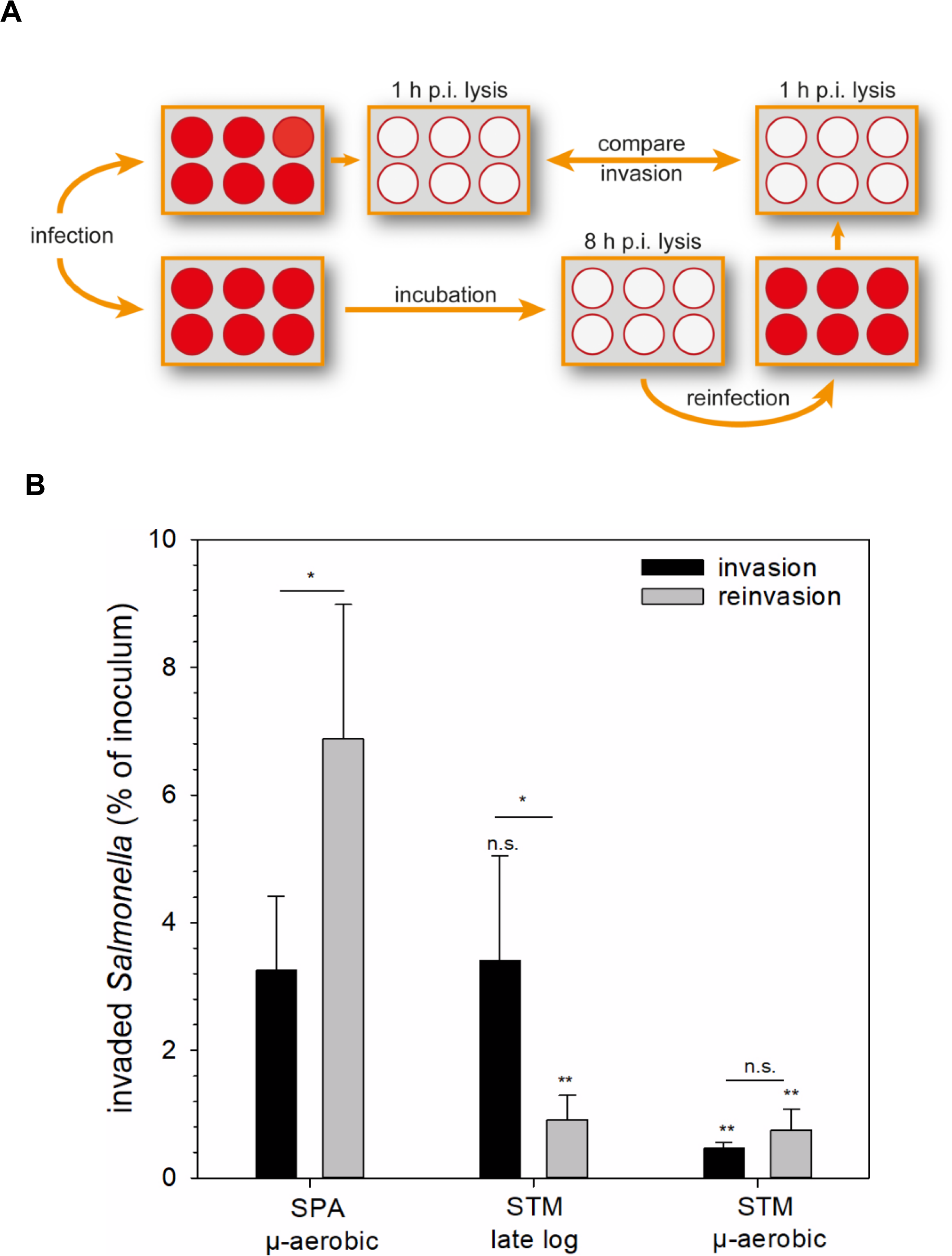
SPA shows higher invasion after release out of infected cells. **(A)** HeLa cells were infected with STM WT or SPA WT at MOI 5, and invasion by each strain was determined by gentamicin protection assays. At 8 h p.i., one portion of infected cells was lysed by hypotonic conditions, and the released bacteria were used to reinfect naive HeLa cells. (**B**) Intracellular CFU counts were determined 1 h p.i. after initial infection (invasion, black bars), or 1 h p.i. after infection with *Salmonella* released from host cells at 8 h p.i. (reinvasion, grey bars). Depicted are means and SD of three biological replicates, each performed in triplicate. For statistical analysis, Student’s t-test was performed. Significance is indicated as n.s., not significant; *, p < 0.05; **, p < 0,01; ***, p < 0.001.

We determined similar levels of invasion by SPA after microaerobic culture and STM after aerobic subculture (3.25% and 3.41% of inoculum, respectively), while STM after microaerobic subculture invaded less efficiently (0.47%). Interestingly, 6.88% of the SPA released from HeLa cells at 8 h p.i. were capable of reinvasion of naïve HeLa cells. In contrast, only 0.91% of released STM were able to reinvade, and reinvasion of STM after microaerobic culture was lower with 0.75%. These data indicate that higher expression of SPI-1 genes and bacterial motility by intracellular SPA results in increased invasiveness of SPA following exit from host cells (Fig. 9B).

## DISCUSSION

Despite high genetic similarity, typhoid and non-typhoidal *Salmonella* serovars differ hugely in their host specificity, the immune response they elicit in humans, and the clinical manifestations they cause (56). Previously, we showed differences in host cell invasion and SPI-1 expression between STM and SPA, following extracellular growth in rich medium LB under aerobic conditions (20) and demonstrated that elevated physiological temperature affects differently their motility and host cell entry (22, 51).

In this work, we focused on their intracellular phase of infection and have taken an unbiased deep RNA-Seq approach to get a panoramic view of the intracellular STM and SPA transcriptomic architecture, and to identify differences in the activity of their genes in the intracellular milieu. Following transition from extracellular to the intravacuolar environment, *Salmonella* undergoes extensive adaptation and global modulation of gene expression to respond to the intracellular environment, however the nutritional contents of the SCV and the metabolic pathways required during intracellular growth are largely undefined. Amino acids and purines seem to be deficient since previous studies have shown that many auxotrophic *Salmonella* mutants are unable to grow intracellularly and are severely attenuated for virulence in mice (57–60). These observations suggest that intracellular *Salmonella* depend on *de novo* synthesis of metabolic precursors.

Recently, Liu and colleagues have applied a proteomic approach to determine the expression of STM proteins within HeLa cells at 6 (61) and 18 h (27) p.i. Also, the STM transcriptome inside macrophages (31, 32) and epithelial cells (26) has been reported previously. Nevertheless, a global transcriptomic comparison between SPA and STM during intracellular infection has not been published thus far.

Previous efforts to identify changes in gene expression using quantitative proteomic approach (LC-MS/MS analyses) have revealed that among about 3,000 detected proteins, the expression of 100 *Salmonella* proteins significantly changed (61 were upregulated and 39 proteins were downregulated) during the transition of STM from the extracellular to the intracellular niche of HeLa cells, at 6 h p.i. (61). A microarray hybridization-based technique has identified 605 upregulated and 616 downregulated STM genes at 6 h p.i. of HeLa cells (26). Our current transcriptomic analysis identified 830 differentially expressed STM genes (365 upregulated and 465 downregulated) that were altered at 8 h p.i., relative to their extracellular expression. Similar numbers of DEGs were also detected for SPA, which presented 322 upregulated and 579 downregulated genes, during epithelial cell infection.

In agreement with these previous analyses, we observed substantial downregulation in the expression of genes associated with SPI-1, Type I fimbriae, flagella and chemotaxis systems in STM, but significant upregulation of STM genes involved in metal ions uptake, purines and biotin synthesis. In fact, nearly all known *Salmonella* iron-, zinc-, or manganese-responsive pathways were found to be intracellularly induced. Since the SCV is characterized by low levels of magnesium, manganese, iron and zinc metals (62) and given the pivotal role of these metal ions in *Salmonella* infection (63, 64), upregulation of iron, zinc and manganese acquisition system genes is not surprising. Nonetheless, these results highlight the importance of biotin and purines to the intracellular phase of *Salmonella* during infection. This notion is consistent with (i) SCVs of both macrophage and epithelial cells are limited for purines and pyrimidines (65); (ii) the avirulent phenotype of purine auxotrophic mutants of *S*. Typhimurium and *S*. Dublin in mice (59); and (iii) the reported contribution of biotin sulfoxide reductase to oxidative stress tolerance and virulence of *Salmonella* in mice (66).

Interestingly, in contrast to the proteomics (61) and the microarray-based transcriptome analyses (26), we were also able to identify significant upregulation of SPI-2 and PhoPQ regulon genes both in STM and SPA. These differences may reflect the time point differences (8 h p.i. in our analysis vs. 6 h p.i. in previous analyses), or the superior sensitivity of the current RNA-Seq techniques.

While the change in the expression of metal transport systems, purines, biotin, PhoPQ and SPI-2 regulons was common to both SPA and STM, intriguing differences in the expression profile of SPI-1, motility and chemotaxis, and carbon utilization pathways (mainly citrate, galactonate and ethanolamine) were identified between intracellular SPA and STM. Ethanolamine is a ubiquitous molecule within vertebrate hosts and serves as a carbon and nitrogen source for bacteria in the intestine, as well as within epithelial cells (67). *Salmonella* can utilize ethanolamine, as an exclusive source of carbon and nitrogen, generating acetaldehyde and ammonia. The ammonia can then provide a cellular supply of reduced nitrogen, while the acetaldehyde can be converted into the metabolically useful compound acetyl-CoA (68). Our data indicated much higher expression of multiple ethanolamine utilization genes by STM vs. SPA during host cell infection and more efficient utilization *in-vitro*. Recently, it was shown that ethanolamine metabolism in the intestine enables STM to establish infection and that EutR, the transcription factor of the ethanolamine cluster in *Salmonella* directly activates the expression of SPI-2 in the intracellular environment (69, 70).

The sugar acid galactonate is another example of an available carbon and energy source that is produced in the mammalian gut. The gut microbiota releases sugar acids including galactonate from ingested polysaccharides present in nutrients by the host, or from simple sugars as catabolic intermediates of metabolism (71). These results may suggest that STM utilizes galactonate, citrate and ethanolamine more effectively than SPA as available carbon sources, and that STM may better coordinate ethanolamine metabolism and virulence via EutR during infection. This notion is consistent with our *in-vitro* data showing that while STM is able to utilize ethanolamine and citrate as a sole carbon source, SPA cannot grow or grows much slower on these sources. Previous studies have shown that deficiency for citrate lyase activity (*citF*) leads to significant reduction in STM survival inside macrophages (72) and that an STM mutant that cannot utilize citrate is highly attenuated in the mouse (73). Similarly, ethanolamine is released by host tissues during inflammation and a study by Thiennimitr and colleagues in mice has shown that ethanolamine metabolism provides a growth advantage to STM during intestinal colonization (70). Since SPA provokes a noninflammatory disease, it is possible that SPA has lost the ability to utilize ethanolamine as it provides no advantage during noninflammatory infection.

It is well established that the expression of the flagella and chemotaxis are highly repressed by intracellular STM (27, 31, 61, 74), supporting a nonmotile state of intracellular STM. In agreement with these reports, we have also observed extensive repression of the entire motility-chemotaxis regulon in intracellular STM, however we found significantly higher expression of this regulon in intracellular SPA. Similar observations were also obtained for the evolutionary related T3SS-1 genes. Most salmonellae are motile and express peritrichous flagella around its membrane. Like T3SS-1, *Salmonella* motility is playing an important role in host colonization and virulence (75, 76). Previous reports have shown a regulatory association between T3SS-1 and the flagella in STM (77). Subsequent to STM entry into host cells, the intracellular expression of T3SS-1 (27, 31, 78) as well as flagella production and chemotaxis (27, 31, 61, 79) genes were shown to be highly repressed, possibly to prevent inflammatory response by the NAIP/NLRC4 inflammasome (80). Therefore, it was unexpected to find that the flagella regulon is readily expressed during SPA intracellular infection and that it maintains an intracellular motility during epithelial cell infection.

Intracellular motility is long known in bacterial pathogens such as *Listeria monocytogenes, Shigella flexneri*, *Rickettsia* spp., and *Burkholderia* spp., that all exhibit intracellular actin-based motility to spread from cell to cell (81). Nonetheless, to the best of our knowledge, this is the first report of a flagellum-dependent intracellular motility of any *Salmonella* serovar in living host cells. While we did not find evidence for intercellular spread, it is possible that this mechanism facilitates escape of SPA from the SCV to the cytoplasm. Previous studies have shown that a small subpopulation of hyper-replicating STM resides within the cytosol of epithelial cells and serves as a reservoir for dissemination. These motile bacteria retain both SPI-1 and flagella expression, but present a low expression of SPI-2 genes, and therefore are primed for invasion (82). This expression profile is largely similar to the intracellular profile of SPA. Indeed, we showed that intracellular SPA possesses higher ability to reinfect non-phagocytic cells than intracellular STM. We propose that elevated expression of motility-chemotaxis and SPI-1 genes in addition to flagella-mediated motility prime SPA towards a new cycle of host cell infection, supporting systemic dissemination of this pathogen in the human body. These phenotypes may contribute to the invasive nature of paratyphoid fever and the ability of SPA to disseminate beyond the intestinal lamina propria of the human host. Future work has to reveal which cells or tissues of infected host organisms are affected by intracellular motility of SPA. However, such analyses are hampered by the lack of animal models for infection. Infection models with human organoids (83), for examples based on gall bladder epithelial organoids (84) as a tissue type important for SPA persistence and dissemination, may offer new perspectives for analyses of cellular interactions.

## MATERIAL AND METHODS

### Bacterial strains, growth conditions and data sets

Bacterial strains utilized in this study are listed in **Table S6**. *S*. Typhimurium 14028 S (85) and *S*. Paratyphi A strain 45157, an epidemic strain, responsible for a paratyphoid outbreak in Nepal (9) were used as the wild-type strains. Bacterial cultures were routinely maintained in Luria-Bertani (LB; BD Difco) liquid medium at 37°C. An M9 medium [33.7 mM Na_2_HPO_4_, 22 mM KH_2_PO_4_, 8.5 mM NaCl, 18.7 mM NH_4_Cl, 12.1% Trizma base, pH=7.4] supplemented with 2 mM MgSO_4_, 0.3 mM CaCl_2_, 3.7 µM thiamine, 4.1 µM biotin, 0.134 mM EDTA, 31 µM FeCl_3_, 6.2 µM ZnCl_2_, 7.6 nM CuCl_2_, 4.2 nM CoCl_2_, 1.62 µM H_3_BO_3_, 8.1 nM MnCl_2_, with or without 0.1% Casamino acids, and with one of the following carbon sources glucose (11 mM), ethanolamine hydrochloride (51 mM) or sodium citrate (34 mM) was used as a minimal growth medium.

For HeLa cell infections, *Salmonella* cultures were grown to the stationary phase under microaerobic conditions (see below). The final dataset for RNA-seq included 12 samples resulting from three independent growth in LB or infection experiments that were used for RNA extraction: (i) *S*. Typhimurium 14028 S bacterium grown in LB medium (hereafter named STM_m) (ii) *S*. Paratyphi A strain 45157 grown in LB medium (hereafter named SPA_m) (iii) Typhimurium 14028 S bacterium infected Hela-cells (hereafter named STM_i) and (iv) *S*. Paratyphi A strain 45157 bacterium in Hela-cells (hereafter named SPA_i).

### Infection of HeLa cells

Human epithelial HeLa (ATCC CCL-2) cells were purchased from the American Type Culture Collection and were cultured in a high-glucose (4.5 g/l) DMEM supplemented with 10% FBS, 1 mM pyruvate and 2 mM L-glutamine at 37 °C in a humidified atmosphere with 5% CO_2_. Cells were seeded at 6×10^6^ cells/ml in a onewell plate (Greiner) tissue culture dish 18 h prior to bacterial infection and infected at multiplicity of infection (MOI) of ∼200 (bacteria per cell). For RNA extraction, five plates for *S.* Typhimurium and six plates for *S.* Paratyphi A were used. *Salmonella* strains carrying the pBR-GFP2 plasmid were grown in 2 ml LB containing 20 μg/ml tetracycline at 37 ^°^C with shaking (250 rpm) for 6-8 h, and then diluted 1:75 into 10 ml LB supplemented with 20 μg/ml tetracycline and grown without shaking at 37 ^°^C for 16 h. Bacterial cultures were centrifuged for 5 min at 4,629 × g, at room temperature (RT), and resuspended in 3 ml prewarmed complete DMEM. 3 ml of DMEM suspended Salmonellae (∼2.5×10^9^ CFU) were used to replace the medium of seeded cells and plates were centrifuged for 5 minutes at 142 × g at RT. After centrifugation, 7 ml of prewarmed DMEM were added and plates were incubated at 37 ^°^C under 5% CO_2_ atmosphere. At 2 h post infection (p.i.) the medium was aspirated, cells were washed three times with 10 ml PBS containing CaCl_2_ and MgCl_2_ (PBS +/+), and incubated with 10 ml prewarmed DMEM containing 100 µg/ml gentamicin. After 60 min, the media was replaced with DMEM containing 10 µg/ml gentamicin and cells were further incubated as before. At 8 h p.i., the cells were washed three times with 10 ml PBS +/+ and trypsinized with 2 ml prewarmed trypsin at 37 ^°^C. When the cells were detached, prewarmed DMEM was added to quench the trypsin. The cells were transferred to a 50 ml conical tube, precipitated by 5 min centrifugation at 247 x g and washed in 15 ml PBS+/+. Finally, the cells were resuspended in 1 ml / plate fixation reagent (RNAlater: RNAprotect 350:1), incubated at RT for 20 min, and then stored at 4 °C until sorting.

HeLa cell lines stably transfected for expression of LAMP1-GFP or Lifeact-GFP have been described before (86) and were maintained and infected as described above.

### FACS sorting and RNA extraction

RNA was extracted from three independent infection experiments of HeLa cells with STM strain ATCC 14028S or SPA 45157. As controls, RNA was also extracted from three independent cultures of uninfected cells and from bacterial cultures grown in LB. HeLa cells (infected or uninfected) were fixed, centrifuged at 247 × g for 5 min and resuspended in 3 ml PBS+/+. The cells were filtered through 30 µm pre-separation filter into 5 ml FACS tubes (SARSTEDT) and sorted on a BD-FACSAria IIu (100 µm nozzle; ND filter 2) at a rate of approximately 300 cells/sec. 2-2.5×10^6^ GFP-positive (infected) HeLa cells were collected and centrifuged at 247 × g for 5 min. RNA extraction was performed using Hybrid-R RNA Purification System (GeneAll). RNA samples were treated with RNase-free DNase (Qiagen) according to manufacturer’s protocol, followed by ethanol precipitation and Ribosomal RNA depletion that was performed by Epidemiology Ribo-Zero Gold rRNA Removal Kit (Illumina).

### RNA sequencing

RNAseq was performed at the GeT-PlaGe core facility, INRAE Toulouse. RNA-seq libraries were prepared using the Illumina TruSeq Stranded mRNA Library Prep, without a poly-A mRNA selection, according to the manufacturer’s protocol. Briefly, RNAs were fragmented to generate double-stranded cDNA and adaptors were ligated. 11 cycles of PCR were applied to amplify the libraries and their quality was assessed using a Fragment Analyser. Libraries were quantified by qPCR using the Kapa Library Quantification Kit (Roche). To reach 5 million bacterial reads for each sample, as recommended in (87), RNA-seq reads were generated from three lanes of an Illumina HiSeq3000 and two lanes of an Illumina NOVAseq6000 using a paired-end read length of 2×150 bp. Ribosomal RNA (rRNA) reads were filtered out using sortmerna-2.1b (88) against Silva and Rfam databases. The raw reads data for this study have been deposited in the European Nucleotide Archive (ENA) at EMBL-EBI under accession number PRJEB46495 and as specified in Table S7.

### Genome annotation and quality control

A flowchart summarizing the RNA-seq pipeline and bioinformatic analyses is shown in Fig. S4. Annotation of SPA 45157 and STM 14028S genomes was performed using EugenePP 1.2 (89) with general and targeted evidence to predict coding genes and noncoding features (Figure S1). Functional prediction was produced with InterProscan v5.15-54.0 with GO terms, COG version COG2014 (90), REBASE v708 (91), IslandViewer4 (92) and PHAST (93).

Structural and regulatory non-coding RNA were annotated using RNAspace software v1.2.1 (94) together with the following tools: RNAmmer 1.2 for rRNA (95), tRNAScan-SE 1.23 for tRNA (96) and blastn 2.2.25 against RFAM 10.0 for other ncRNA (97). The gff3 files produced by Eugene and RNA annotation were merged by a semi-manually procedure using BEDTools-2.26.0 (98).

FastQC v0.11.7 was used for quality control of the raw sequence data. Cutadapt 1.8.3 was used to remove Illumina Truseq adaptors and to filter out all reads with length shorter than 36 nucleotides.

### Reads mapping and counting

The remaining, non rRNA reads were mapped with STAR-2.6.0c (99), with default parameters, against a “hybrid” genome composed by the Ensembl GRCh38 human primary assembly (release 92) and the SPA (accession number CP076727) or STM (accession number CP001362 - CP001363) genomes. In addition, in order to analyze bacterial samples, we indexed each bacterial genome separately by decreasing the parameter “genomeSAindexNbases” to 10 for STM and to 9 for SPA, while producing the alignment (.bam) files sorted by coordinates. Samtools flagstat v1.8 was used to obtain mapping statistics. Then we added read groups to trace the origin of each read with AddOrReplaceReadGroups of Picard tools v2.18.2. Next, we merged the bam files that originated from resequencing of the same sample using the MergeSamFiles command of Picard tools. The featureCounts command from the Subread-1.6.0 tool (100) was used to compute fractional counting for all annotated genes.

### Clustering of orthologous genes

Orthologous genes were computed using Roary v.3.11.2 (101) with default parameters (minimum of 95% identity between coding sequences of the same cluster) except for the “-s” option that avoids paralog splitting in different clusters. Based on the orthologous groups, we produced raw count matrices that compare the expression level between orthologous genes of SPA and STM. We also added groups including paralog genes on either SPA or STM genome into the raw count matrix to sum up the raw reads count of each putative paralog gene.

### Differential Gene expression analysis

Normalization and differential analysis were carried out according to the DESeq2 model and package (102) using adjusted p-value with FDR correction and the Wald test to infer the probability value. Genes that had too few reads (the sum of reads for all the samples was less than 10) were discarded. To compare DEGs between SPA and STM, we created a count matrix composed of the shared protein-coding and ncRNA genes between these two serovars. The heatmaps were created by applying DESeq2 variance stabilizing transformations (the vst command) on the counts followed by subtracting the mean of the gene across all samples. The heatmaps were drawn using the ComplexHeatmap R package (103).

### Reverse transcription and quantitative real-time PCR (qRT-PCR)

500 ng of RNA was reverse-transcribed using qScript cDNA synthesis kit (Quanta-bio). Real-time PCR was performed as previously described (20) on a StepOnePlus Real-Time PCR System (Applied Biosystems). Relative quantity of transcripts was calculated as 2^-ΔΔCt^, using the *rpoD* gene as an endogenous normalization control.

### Western Blotting

HeLa cells were infected with *Salmonella* strains carrying pBR-GFP2 plasmid and pWSK29 carrying HA-tagged alleles of *sopE2* and *sopB*. The infection procedure was performed as above until the fixation step. Cell fixation was performed by incubation with 5 ml of 4% formaldehyde for 15 min under gentle shaking. Five ml of PBS +/+ were added and the cells were centrifuged for 5 min at 2,700 × g. Subsequently, the cells were washed three times with PBS +/+ and finally resuspended in PBS +/+ containing 1% BSA. The cells were stored at 4 °C until sorting. 1×10^6^ GFP-positive cells were centrifuged for 5 min at 247 × g, and resuspended in 100 μl sample buffer. Western blot of the infected cells was performed as previously described (104) using the anti-2HA (Abcam ab18181; 1:1,000), anti-FliC (Abcam ab93713; 1:200,000) and anti-DnaK (Abcam ab69617; 1:10,000). Proteins were quantified relative to DnaK using ImageJ.

### Host cell infection and live-cell imaging

HeLa cells stably transfected with LAMP1-GFP were seeded in surface-treated 8-well slides (ibidi) 24 or 48 h prior to infection to reach ∼80% confluency (∼80,000 cells) and were then used for infection. STM strains for infection experiments were subcultured from an overnight culture (1:31) in fresh LB medium and grown for 3.5 h at 37°C. For infection, SPA strains were grown for 8 h under aerobic conditions, subcultured (1:100) in fresh LB medium and stationary-phase subcultures were grown for 16 h under microaerophilic conditions as described in (21). Bacteria were adjusted to an optical density of 0.2 at 600 nm in PBS and used for infection with the respective MOI (between 25 and 100, dependent on strain). Bacteria were centrifuged onto the cells for 5 min at 500 × g to synchronize infection and the infection was allowed to proceed for 25 min. After washing thrice with PBS, cells were incubated in medium containing 100 µg × ml^-1^ of gentamicin to kill extracellular bacteria. Afterwards, the cells were maintained in medium supplemented with 10 µg × ml^-1^ gentamicin for the ongoing experiment. Live-cell imaging was performed with Cell Observer Spinning Disc microscope (Zeiss) equipped with a Yokogawa Spinning Disc Unit CSU-X1a5000, an incubation chamber, 63× objective (α-Plan-Apochromat, NA 1.4), two ORCA Flash 4.0 V3 cameras (Hamamatsu) and appropriate filters for the respective fluorescence proteins.

### Immunostaining of infected cells

HeLa cells stably transfected with LAMP1-GFP were seeded in surface-treated 24-well plates (TPP) on coverslips 24 h or 48 h prior infection to reach ∼80% confluency (∼180,000 cells) and were used for infection. Cultivation of bacteria and infection of HeLa cells was carried out as described above. After the desired incubation time the cells were washed thrice with PBS and fixed with 3% PFA in PBS for 15 min at RT, followed by immunostaining. Fixed cells were washed thrice with PBS and incubated in blocking solution (2% goat serum, 2% bovine serum albumin, 0.1% saponin, 0.2 % Triton X-100) for 30 min. Cells were stained 1 h at RT with primary antibodies against *Salmonella* (H:i, BD Difco 228241; H:a, Sifin TR1401; O:5, Sifin TR5303). After washing thrice with PBS, cells were incubated with the appropriate secondary antibodies for 1 h at RT. Coverslips were mounted with Fluoroshield (Sigma) and sealed with Entellan (Merck). Microscopy of fixed samples was either performed using a Cell Observer Spinning Disc microscope (Zeiss) as described above, or using a Leica SP5 confocal laser-scanning microscope using 100× objective (HCX PL APO CS, NA 1.4-0.7) and polychroic mirror TD 488/543/633.

### Reinvasion assays

HeLa cells were seeded in surface-treated 6-well plates (TPP) 24 h or 48 h prior to infection to reach ∼100% confluency (1×10^6^ cells) on the day of infection. Infection cultures of STM and SPA strains were grown as either 3.5 h aerobic or 16 h microaerobic subcultures and used for infection as described above. At 1 h p.i., cells were washed thrice with PBS and lysed using 0.1% Triton X-100. CFU were determined by plating serial dilutions of lysates and inoculum on Mueller-Hinton II (MH) agar and incubated overnight at 37 °C. The percentage of internalized bacteria of the inoculum (from subcultures) was calculated. 8 h p.i., cells were washed twice with PBS, once with ice-cold ddH_2_O, and lysed using ddH_2_O. Bacteria liberated from host cells by hypotonic lysis were used to infect naïve HeLa cells, and 1 h p.i. cells were washed and lysed and CFU were determined as described above. The percentage of internalized bacteria of the inoculum (from liberated bacteria) was calculated.

## ACKNOWLEDGMENTS

This work was supported by the Infect-Era program, SalHostTrop project (“Understanding the Human-Restricted Host Tropism of Typhoidal Salmonella”, 2016-2020) and by the France Génomique National infrastructure, funded as part of “Investissement d’avenir” program managed by Agence Nationale pour la Recherche (ANR-10-INBS-09 contract). The work at the Gal-Mor laboratory was supported by grant numbers: 2616/18 from the joint ISF-Broad Institute program; 3-12435 from Infect-Era /Chief Scientist Ministry of Health; I-41-416.6-2018 from the German-Israeli Foundation for Scientific Research and Development (GIF, awarded to OGM and MH); and A128055 from the Research Cooperation Lower Saxony – Israel (The Volkswagen Foundation, awarded to OGM, MH and GG). The funders had no role in study design, data collection, and interpretation, or the decision to submit the work for publication. We thank C. Gaspin for providing annotation of ncRNA genes of the STM and SPA genomes.

## SUPPLEMENTARY MATERIALS

**Fig. S1.**
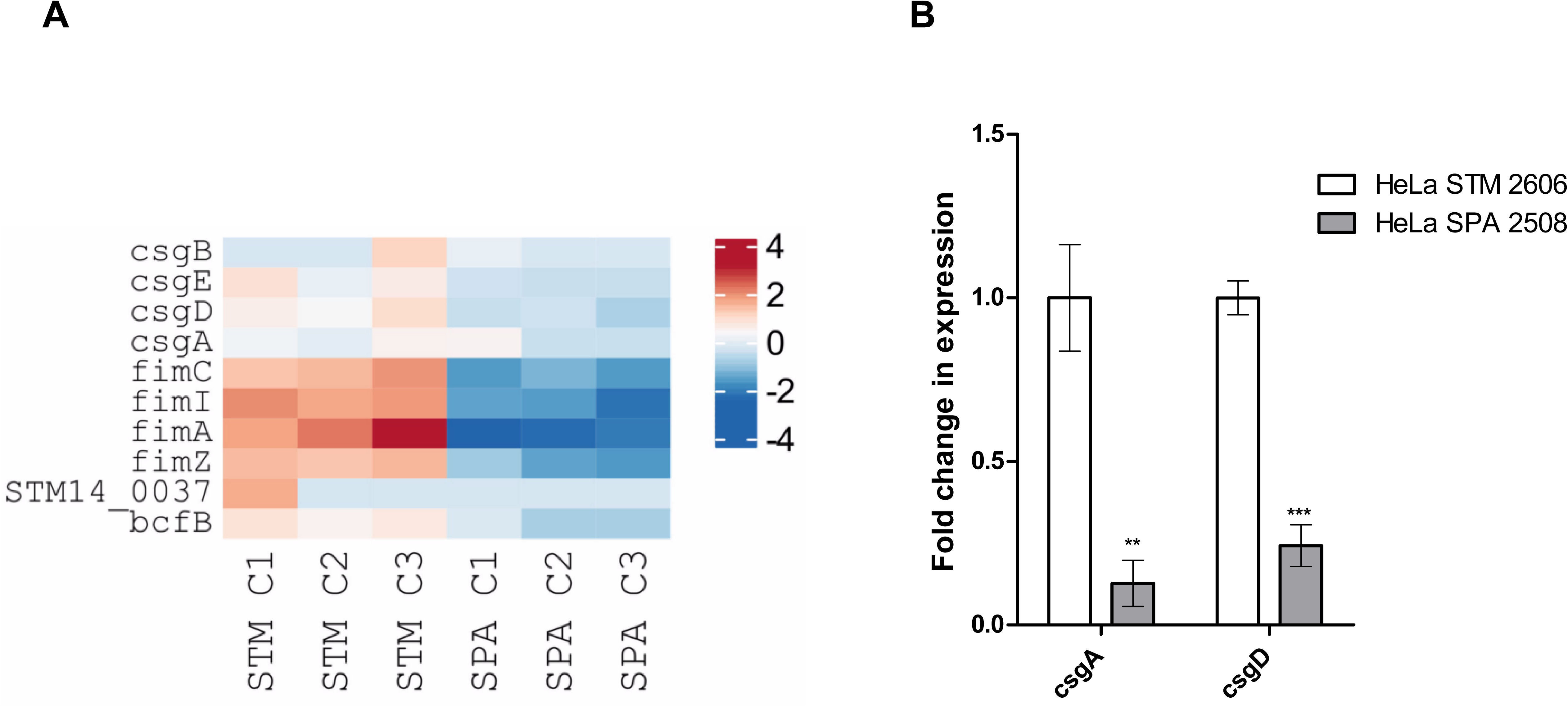
Expression of fimbriae genes in intracellular STM vs. SPA. (**A**) Heat map of RNA-Seq results showing the relative transcription of genes encoding curli (Csg), Fim, or Bcf fimbriae in three independent HeLa cells infections with STM and SPA. (**B**) The x-fold change in the expression of *csgA* and *csgD* in intracellular SPA relative to their expression in intracellular STM was analyzed by qRT-PCR. The results show the means of 4-8 independent reactions and the error bars indicate the SEM.

**Fig. S2.**
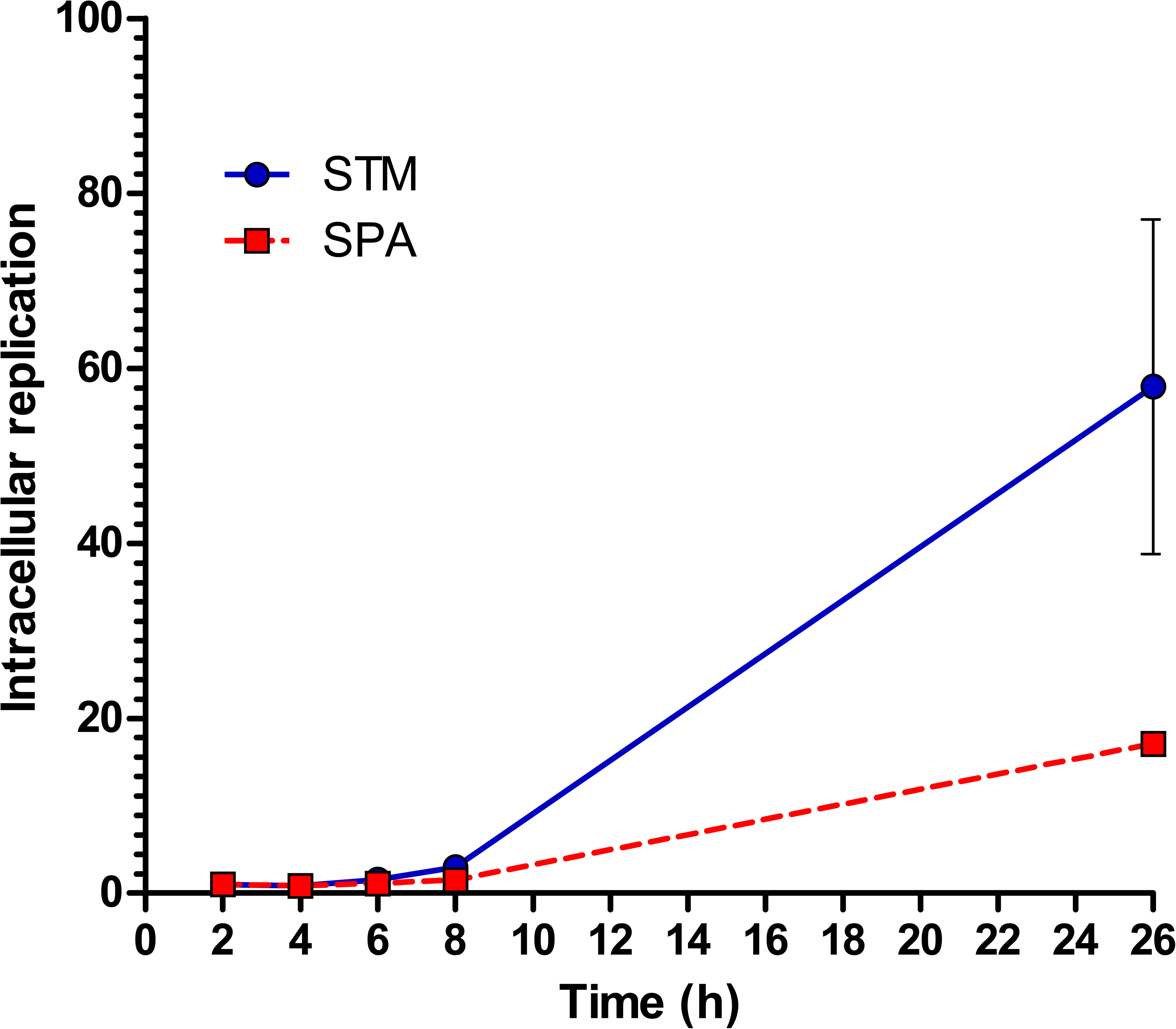
Intracellular replication of SPA and STM in HeLa cells. Intracellular growth of STM vs. SPA in HeLa cells at 2, 4, 6, 8 and 24 h p.i. Bacterial replication was determined by the gentamicin protection assay. Replication was calculated as the ratio between the intracellular bacteria (CFU) recovered at each time point and the number of CFU at 2 p.i. The means and the SEM of four independent infections are shown.

**Fig. S3.**
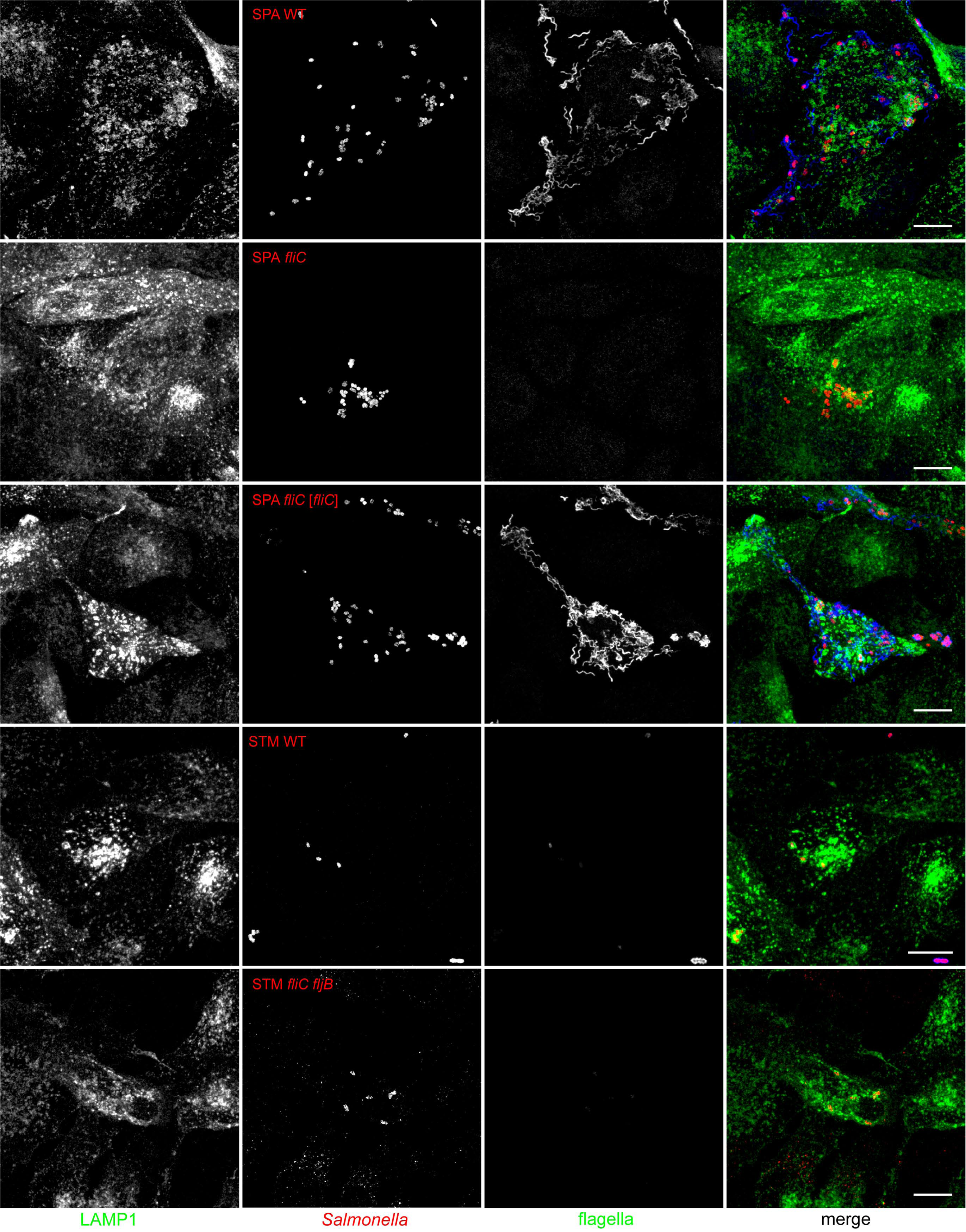
Formation of flagella by SPA within the host cell cytoplasm. HeLa cells stably expressing LAMP1-GFP (green) were infected with SPA WT, SPA *fliC*, SPA *fliC* [*fliC*], STM WT, or STM *fliC fljB* as indicated, each constitutively expressing mCherry (red). Infected cells were fixed 8 h p.i. and subsequently immunolabeled with serovar-specific antibody against FliC (blue). Scale bars: 10 µm.

**Fig. S4.**
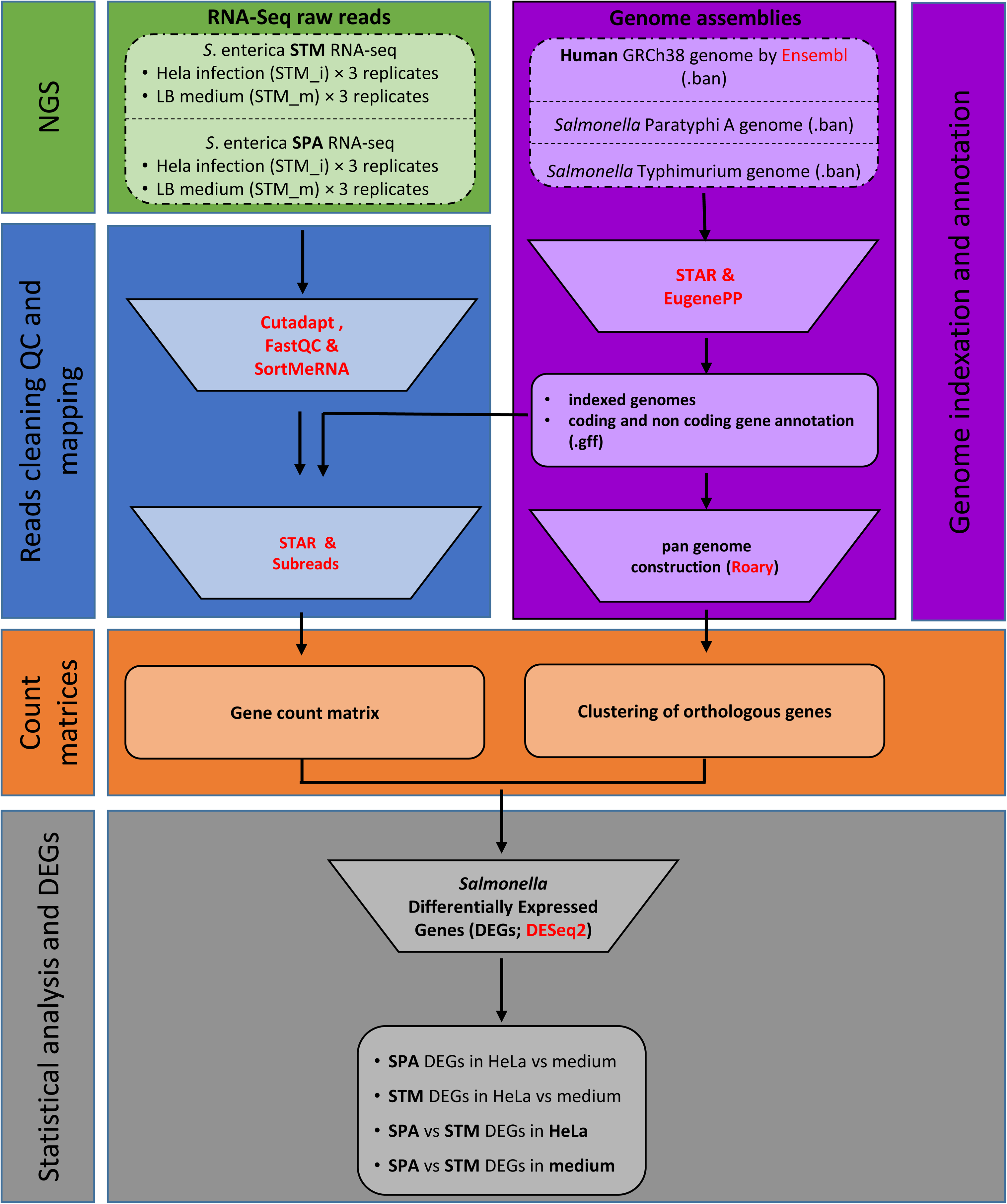
A flowchart of RNA-seq and bioinformatic analyses. The different steps and main tools used for the bioinformatic analyses illustrated. RNA extracted from STM and SPA grown in culture and from infected cells was subjected to NGS on Illumina sequencing. RNA-seq reads were cleaned and quality controlled using FastQC. Cutadapt was used to remove Illumina Truseq adaptors and to filter out all reads shorter than 36 nucleotides. SortMeRNA was used to filter out ribosomal RNA reads from raw data. Human and bacterial genomes were merged into “Hybrid genomes” and indexed with STAR. Bacterial genomes were reannotated using EugenePP. STAR was used to align reads to reference genomes and sort resulting bam files by coordinates. To compute fractional counting for all annotated coding and non-coding genes the Subread tool with parameters “-s 2 (reversely stranded), -p (count fragments instead of reads) -M, -O, --fraction” was used. Clustering of *Salmonella* homologous genes was computed using Roary. Normalization and differential expressed genes analysis were carried out according to the DESeq2 model and package using adjusted P-value and the Wald test to infer the probability value. To obtain DEGs between STM and SPA a count matrix composed of the shared and coding genes and ncRNAs between these two serovars was created.

## TABLES

**Table S1.** The transcriptome of STM grown in LB and intracellularly. The RNA-seq reads obtained from STM grown on LB or extracted from HeLa cells infected with STM were mapped to the STM 12048S genome. This table shows the normalized (TPM) featurecount counts of reads assigned to the STM genes. This table also lists the genes that were found expressed in LB only, intracellularly only, and at both conditions.

**Table S2.** DEGs of intracellular STM vs. LB.

**Table S3.** The transcriptome of SPA grown in LB or intracellular. The RNA-seq reads obtained from SPA grown on LB or extracted from HeLa cells infected with SPA were mapped to the SPA 45157 genome. This table shows the normalized (TPM) featurecount counts of reads assigned to the SPA genes. This table also lists the genes that were found expressed in LB only, intracellular only, and under both conditions.

**Table S4.** DEGs of intracellular SPA vs. LB.

**Table S5.** DEGs between intracellular STM and intracellular SPA.

**Table S6.** Bacterial strains and primers utilized in this study.

**Table S7.** RNA-Seq metadata and accession numbers of the 12 samples analyzed in this study.

## MOVIES

**Movie 1. Live-cell imaging of STM and SPA strains reveals intracellular motility of SPA.** To investigate intracellular motility, HeLa cells stably expressing LAMP1-GFP (green) or LifeAct-GFP (green) as indicated were infected with SPA WT, SPA *ssaR*, SPA *fliC*, or STM ATCC 14028s WT, each harboring pFPV-mCherry for constitutive expression of mCherry (red). Individual clips of about 30 seconds are shown successively. Time stamps in upper left corner indicate sec.ms. Scale bars: 10 µm.

**Movie 2. Real time imaging of intracellular motility of SPA WT.** HeLa cells stably expressing LifeAct-GFP (green) were infected with SPA WT expression mCherry (red) and imaged at a frame rate of ∼40 frames per second. Time stamps in upper left corner indicate sec.ms. Scale bar: 10 µm.

**Movie 3. Real time imaging of intracellular motility of SPA WT.** HeLa cells stably expressing LifeAct-GFP (green) were infected with SPA WT expression mCherry (red) and imaged at a frame rate of ∼40 frames per second. Time stamps in upper left corner indicate sec.ms. Scale bar: 10 µm.

**Movie 4. Live-cell imaging of further intracellular STM and SPA strains.** HeLa cells stably expressing LAMP1-GFP (green) were infected with STM SL1344 WT, or various isolates of SPA as indicated in the lower left corner of movie sections. All strains harbored pFPV-mCherry for constitutive expression of mCherry (red) Time stamps in upper left corner indicate sec.ms. Scale bars: 10 µm.

