## Supplementary material for "Intracellular *Salmonella* Paratyphi A is motile and differs in the expression of flagella-chemotaxis, SPI-1 and carbon utilization pathways in comparison to Intracellular *S*. Typhimurium": Table S6

**Table S6. Bacterial strains and plasmids used in the study**

| **Strains** | **Description** | **Reference or source** |
| --- | --- | --- |
| *S*. Typhimurium SL1344 | wild type Sm^r^ *xyl hisG rpsL* | SGSC |
| *S.* Typhimurium 14028s | wild type | SGSC |
| *S.* Typhimurium MvP1760 | Δ*fliC*::FRT Δ*fljB*::aph | 1 |
| *S.* Typhimurium P2D6 | *ssaV*::mTn5 | 2 |
| *S.* Paratyphi A (SPA) 45157 | wild type, clinical isolate, 2009 Nepal outbreak | 3 |
| SPA 45157 *ssaR* | Δ*ssaR*::FRT | This study |
| SPA 45157 *fliC* | Δ*fliC*::FRT | This study |
| *S.* Paratyphi A 113498 | wild type, 2008 clinical isolate, traveler from Sri Lanka | 3 |
| *S.* Paratyphi A 93223 | wild type, 2004 clinical isolate, traveler from Romania | 3 |
| *S.* Paratyphi A 105493 | wild type, 2006 clinical isolate, traveler from Thailand and Nepal | 3 |
| *S.* Paratyphi A 83698 | wild type, 2003 clinical isolate, traveler from India, | 3 |
| *S.* Paratyphi A SARB 42 | wild type | 4 |
| *S.* Paratyphi A SARB MZ763 | wild type | 4 |
| **Plasmids** | | |
| pBR::GFP2 | GFP in pBR322, tet^r^ | This study |
| pWSK29::SopB-2HA | SopB from STM SL1344 and SPA 45157, respectively, with C-terminal two-hemagglutinin tag (2HA) in pWSK29, amp^r^ | 5 |
| pWSK29::SopE2-2HA | SopE2 from STM SL1344 and SPA 45157, respectively, with C-terminal two-hemagglutinin tag (2HA) in pWSK29, amp^r^ | 6 |
| pFPV-mCherry |  |  |
| p5141 | SPA fliC PEM7::tagRFP-T | This study |

SGSC – Salmonella genetic Stock Center the University of Calgary.

1 Fulde, M., Sommer, F., Chassaing, B., van Vorst K., Dupont, A., *et al*. Neonatal selection by Toll-like receptor 5 influences long-term gut microbiota composition. Nature (2018) 560, 489–493

2 Shea, J. E., Hensel, M., Gleeson, C., Holden, D.W. Identification of a virulence locus encoding a second type III secretion system in *Salmonella* typhimurium. Proceedings of the National Academy of Sciences (1996) 93 (6) 2593-2597.

3 Gal-Mor, O., Suez, J., Elhadad, D., Porwollik, S., Leshem, E., Valinsky, L., *et al*. Molecular and cellular characterization of a Salmonella enterica serovar Paratyphi A outbreak strain and the human immune response to infection. Clinical and vaccine immunology: CVI (2011)19(2):146-156

4 Boyd, E.F., Wang, F-S., Beltran, P., Plock, S.A., Nelson, K., *et al.* *Salmonella* reference collection B (SARB): strains of 37 serovars of subspecies I. Journal of General Microbiology (1993)139, 1125-1132

5 Elhadad, D., Desai, P., Rahav, G., McClelland, M., Gal-Mor, O. Flagellin Is Required for Host Cell Invasion and Normal Salmonella Pathogenicity Island 1 Expression by Salmonella enterica Serovar Paratyphi A. Infect Immun. (2015) 83(9):3355-68.

6 Elhadad, D., Desai, P., Grassl, G.A., McClelland, M., Rahav, G., Gal-Mor, O. Differences in Host Cell Invasion and Salmonella Pathogenicity Island 1 Expression between Salmonella enterica Serovar Paratyphi A and Nontyphoidal S. Typhimurium. Infect Immun. (2016) 24;84(4):1150-1165.

**Real-Time PCR primers used in the study**

| Primer | Sequence 5’-3’ |
| --- | --- |
| rpoD-F | GGTCTGACCATCGAACAGGTG |
| rpoD-R | ATCAGACCGATGTTGCCTTC |
| uhpT-F | TCTTCGCTGATCTCTTCGCC |
| uhpT-R | TGGCTTTATCGGTCTGCGTT |
| citX-F | TACTCTTCTGCCAGCGTGTG |
| citX-R | GGTCTCCTTTACCGTGGTGG |
| citA-F | GCTGATAGCGGAAGTCGTCA |
| citA-R | ATGACGCTCAGGCCATTGAA |
| eutP-F | GTTTGCCGCGTGGACATAAA |
| eutP-R | CCCGTTGGTATCACGCCTTA |
| eutQ-F | TTTCGCTTTTACACGCCTGC |
| eutQ-R | ACTGGGCTTCACGATTACCG |
| eutG-F | CGATTCTTGATGCTGCCGTG |
| eutG-R | CCTCAATCGCATGCGTCAAC |
| hilA-F | ATATGCCGTTCTGGTCATCC |
| hilA-R | GCCCTGTCCGTACAGTGTTTC |
| hilD-F | GAGATACCGACGCAACGAC |
| hilD-R | CTGCGCTTTCTCTGTGGG |
| invH-F | AACGCTGATAATTCCGCATC |
| invH-R | CGGTCATGAGTTGCTCTTCAT |
| sopB-F | CGGTCATGAGTTGCTCTTCATC |
| sopB-R | GTGGGCAAAAATACGGAAG |
| sopD-F | GACCTGGCACCCGGAATTAC |
| sopD-R | AGTCCTGCCATTCGACAAGC |
| spaO-F | GGCAGTTCAGGAAGATGCTC |
| spaO-R | ACAGAGCGACCGTTTGAGTTG |
| fliC-F | AAGAGAGGACGTTTTGCGG |
| fliC-R | CGAAGATTCCGACTACGCG |
| fliZ-F | AAACATTTCCCACGATCTGC |
| fliZ-R | CGGTAAAGGGGGATTTCTG |
| flgM-F | CCTTTGAAACCCGTTAGCAC |
| flgM-R | GCCGTTTTTAATGCTTCGAC |
| flgF-F | CAGTTGGACTACACCTCCCG |
| flgF-R | CGGGTATATCCTTCAGCGCC |
| cheA-F | AATGGAAAACCTGCTGGATG |
| cheA-R | CTTAGTGCCGCGGTTTCTAC |
| cheB-F | GCACATGTCGGATAGCTTC |
| cheB-R | AGATGATCGCCGAAAAAGTG |
| csgA-F | GTTGTTGCCAAAACCAACCT |
| csgA-R | TTTCAGAAACAATGCCACCA |
| csgD-F | CTGCATAATATTCAACGTTCTCTGG |
| csgD-R | GCCAGTTTTCAATTTCACGGTAG |
